# Phase separation of protein kinase A regulatory subunits is driven by similar inter- and intra-molecular interactions involving the inhibitory segment

**DOI:** 10.1101/2025.08.07.669161

**Authors:** Malyasree Giri, Christopher S. Brasnett, Kübra F. Eroglu, Julie Maibøll Kaasen, Alain A.M. Andre, Siewert-Jan Marrink, Frans A.A. Mulder, Magnus Kjaergaard

## Abstract

Phase separation of biological macromolecules has emerged as a widespread mechanism for cellular organization, governing the spatiotemporal regulation of biochemical processes including signaling pathways. The regulatory subunit of the cAMP-dependent protein kinase A (PKA) RIα phase separates following cAMP-induced kinase activation, thereby restricting and enhancing PKA activity in space and time. The N-terminal dimerization/docking (D/D) domain and the linker region of RIα have been identified as necessary and sufficient for phase separation, but their interactions at atomic resolution have remained elusive. We report here a structural investigation of the minimal phase-separating region of RIα by nuclear magnetic resonance spectroscopy (NMR) and molecular dynamics (MD) simulations. The intrinsically disordered regions of RIα cause broadening of NMR signals throughout the protein indicative of widespread dynamic interactions. Mutation of arginine residues in the RIα inhibitory sequence leads to a reduction in signal broadening accompanied by loss of phase separation. Independently, binding of RIα to an A-kinase anchoring protein (AKAP) leads to spectral changes that are indicative of a loss of interactions between the D/D domain and the linker region, suggestive of a competition between intramolecular interactions and partner binding. Corroborating the experimental observations, multiscale MD simulations detect pervasive interactions of the folded D/D domain of RIα with the intrinsically disordered linker, and the intermolecular interactions in the condensate mirror those found internally in RIα dimers. Our results support the view that for PKA RIα, homotypic phase separation is underpinned by intermolecular interactions that are similar to intramolecular interactions.

## Introduction

3’,5’-cyclic adenosine monophosphate (cAMP) is a second messenger that transmits the signal from G protein-coupled receptors (GPCRs) on the cell surface to internal pathways, where it activates protein kinase A (PKA) and other downstream targets. The catalytic subunit of PKA recognize a short basic motif^1^ and can phosphorylate a myriad of protein substrates.^2^ Enigmatically, PKA regulates the physiology of organs as different as the heart,^3^ brain^4^ and pancreas^5^ where it regulates diverse processes spanning metabolism, gene expression, and cell growth, all based on the same input signal, an elevation in cAMP.

The PKA holoenzyme is an R2C2 heterotetramer composed of a homodimer of regulatory (R) subunits and two catalytic (C) subunits.^6,7^ Each R subunit binds to one C subunit via a linker region that contains an embedded inhibitory sequence (IS).^8^ The IS blocks catalysis until cAMP binding triggers conformational changes that release active catalytic subunits and allow the linker to become fully disordered.^9^ Humans have four isoforms of the R subunit (RIα, RIβ, RIIα, and RIIβ), where RI subunits are primarily found in the cytosol while RII subunits target the kinase to cellular membranes. The four regulatory subunits have a shared domain structure with an N-terminal dimerization/docking (D/D) domain followed by an intrinsically disordered linker and two cyclic nucleotide binding domains (CNB-A and CNB-B) (Fig. 1A). The R subunits display high sequence conservation in their D/D and CNB domains, whereas the linker regions vary significantly in sequence,^10^ and this variation impacts the dynamics and activation threshold of the holoenzyme.^11^

**Figure 1:**
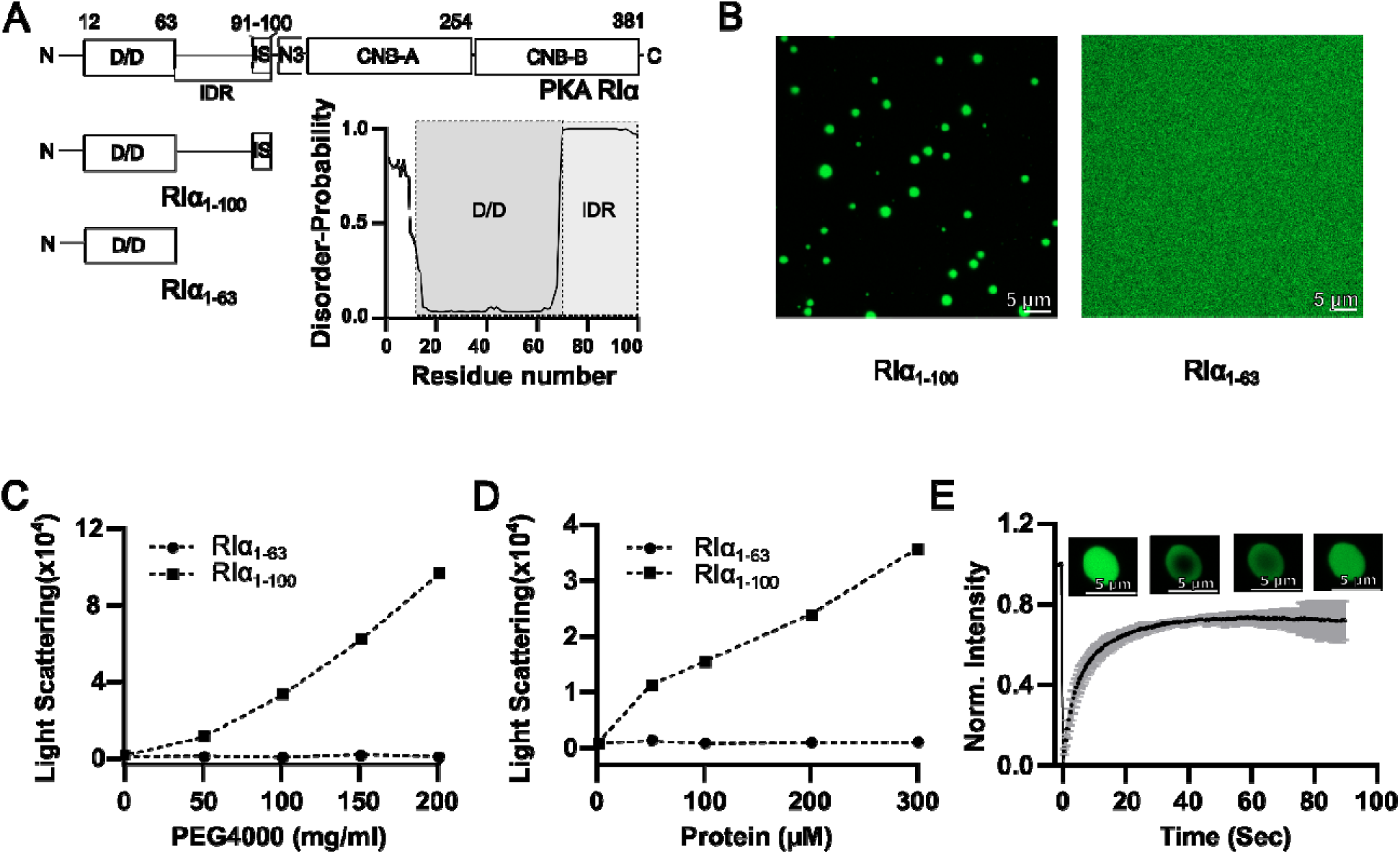
The IDR tail plays a crucial role in RIα phase separation. A) Domain architecture of the full-length protein and the N-terminal folded domain (D/D) with and without IDR tail. The probability of disordered region of RIα_1-100_ was predicted using ODiNPred.^76^ B) Representative confocal images of the droplets formed by 100 µM fluorescent-labeled protein RIα_1-100_ and no droplet formed by 100 µM fluorescent-labeled mutant RIα_1-63_ at 10 mg/ml PEG4000. C) Light scattering of 100 µM RIα_1-63_ and RIα_1-100_ at varying PEG4000 concentration. D) Light scattering of RIα_1-63_ and RIα_1-100_ at 100 mg/ml PEG4000. E) FRAP recovery of liquid droplet formed by 100 µM RIα_1-100_ at 100 mg/ml PEG4000.

The specificity of cAMP signaling is regulated at multiple levels including the cellular distribution of “cAMP sources” and “cAMP sinks”, and anchoring proteins that physically link signaling proteins together. cAMP is produced at adenylyl cyclase point sources following activation of nearby G-protein coupled receptors. Although cAMP is diffusible, its cellular distribution is restricted by phosphodiesterase degradation, which act as local “cAMP sinks”. As a result, the local concentration of cAMP varies at the nanoscale,^12^ and stimulation of the cell will affect PKA differently depending on which protein complex it is in. The localisation of PKA is controlled by a heterogenous family of proteins known as the A-kinase anchoring proteins (AKAPs).^13^ AKAPs target PKA regulatory subunits to specific subcellular sites, restricting PKA signaling to microdomains.^14^ In cAMP-independent phosphorylation events, AKAP physically brings the substrate and enzyme together within the AKAP anchored PKA holoenzyme, and the flexibility within the anchored PKA holoenzyme due to the disordered linker region allows for the precise orientation of the enzyme and substrate. The length and flexibility of the linker region influence the conformational dynamics of the tethered AKAP-PKA complex, enabling it either to adjust kinase accessibility across multiple phosphorylation sites on a single substrate or to direct each catalytic subunit toward distinct nearby substrate proteins within the scaffolded environment.^15^

Recent studies suggest that the RIα subunit further contributes to the compartmentalization of cAMP signaling through phase separation^16^ and that this might be a general mechanism for regulation of kinases.^17,18^ Phase separation has recently emerged as a general compartmentalization mechanism, where macromolecules such as DNA, RNA or proteins form dynamic membraneless organelles. Such structures are refered to as biomolecular condensates and arise due to self-assembly of macromolecules through multivalent interactions. Condensates possess distinct physical properties and can regulate biochemical processes by concentrating some components and excluding others. RIα condensates recruit key signaling components including cAMP, regulatory and catalytic subunits of PKA, PDEs, AKAPs, substrates, and a non-canonical active PKA holoenzyme. Simultaneously, they restrict cAMP diffusion and thereby sustain low cytosolic PKA activity while facilitating highly compartmentalized signaling.^16,19^ This phase separation enhances the efficiency and specificity of PKA activation and works in concert with phosphodiesterases to maintain sharply defined signaling domains. Phase separation of PKA RIα may contribute to pathology in fibrolamellar carcinoma, where the oncogenic DNAJB1-PKA_cat_ fusion protein disrupts normal phase separation behavior of the regulatory subunit PKA RIα and leads to a loss of cAMP compartmentalization and altered PKA signaling. Notably, the loss of RIα condensate formation alone is sufficient to drive increased cell proliferation and transformation in cultured fibroblasts, implicating RIα phase separation as a tumor-suppressive mechanism.^16^

The *in vivo* molecular properties of PKA RIα have been mapped through truncation and mutations: Phase separation is strongest for the full length protein, and removal of either the D/D domain or the interdomain linker distrupts phase separation completely.^16^ The minimal construct that phase separates in cells consists of the N-terminal D/D domain and the linker region of PKA RIα.^16^ Reconstituted holoenzymes consisting of RIα and PKA_cat_ form round liquid-like droplets in vitro in the presence of 100 mg/mL crowder. Addition of cAMP increases the propensity to phase separation, hypothetically because cAMP binding cause the release of the inhibitory sequence and linker region.^16^ The propensity to phase separate has also been investigated in cells by mutagenesis: Mutations in the core of the D/D-domain or in the helical interaction interface preceding CNB-A both disrupt phase separation.^19^ This suggests that RIα dimerization is crucial. At the same time, binding of an AKAP to the D/D domain suppresses phase separation in cells, suggesting a role for this domain beyond dimerization. Likewise, the removal of all charged residues in the linker reduced phase separation. In contrast, mutating the inhibitory sequence to a consensus PKA site increased the propensity to phase separate^19^, possibly because phosphorylation dislodges the disordered linker from its binding site in the catalytic domain. Beyond the inhibitory sequence, the linker has been shown to interact with CNB-A, which allosterically tunes the activation threshold.^20^ A further understanding of this central hub in normal and disease biology is lacking, since high-resolution information about the interactions of the linker between the inhibitory sequence and the D/D domain is currently still missing.

High-resolution information about the drivers of phase separation can be obtained by nuclear magnetic resonance (NMR) spectroscopy and molecular dynamics (MD) simulations. NMR has been particularly informative as an experimental method to study intrinsically disordered proteins even inside highly viscous condensates.^21^ NMR was used to study TDP-43,^22^ hnRNPA2,^23^ CAPRIN1,^24,25^ FUS,^26^ tau^27^ among others, and these proteins all remain disordered in the dense phase. Through an analysis of chemical shifts and relaxation rates, the intermolecular contact interaction hotspots can be identified, such as, for example, the helical regions in the low complexity domain of TDP-43^22^ or the aggregation-prone hexapeptide repeats in tau.^27^ Experimental data can be compared to MD simulations, which in principle allow interactions to be studied at atomic resolution. However, the time- and length-scales required to fully describe the intermolecular interactions driving condensate formation are challenging for atomistic force fields. Instead, the proteins can be represented using a coarse-grained (CG) model, where several atoms are combined into beads. For IDPs, one-bead-per-residue models have proven efficient at capturing compaction and phase diagrams of condensation ^28–31^. Intramolecular interactions drive phase separation to an extent where phase diagrams can be predicted from algorithms trained on single-chain compaction.^32–34^ Alternatively, the Martini coarse-grained force field offers a higher resolution whilst still being computationally efficient. Martini uses a transferable building-block approach, representing 2-4 heavy atoms per bead^35–37^, and has been used to simulate several condensate systems.^38–42^

Here, we describe NMR experiments and multiscale MD simulations to probe the structure, dynamics and interactions of PKA RIα with the aim to elucidate the origin of its propensity to form biomolecular condensates. Phase separation is recapitulated in a minimal construct consisting of the D/D domain and the disordered linker, which forms dynamic condensates in vitro. We detect widespread intra- and intermolecular interactions between the folded domain and the IDR of PKA RIα. These interactions depend on a cluster of arginine residues in the inhibitory sequence and the AKAP binding site in the D/D domain. Our results show how ligand binding to the RIα subunit provides a mechanism to modulate phase separation *in vitro*, a process that may explain observations on PKA activity made *in vivo*.

## MATERIALS AND METHODS

### Plasmid constructs of human PKA RIα

The plasmids for wild type and mutant proteins were prepared by de novo synthesis (Genscript Inc.) and codon optimized for expression in *Escherichia coli* based on the sequence for human PRKAR1A (UNIPROT P10644). The coding regions on were cloned into pET28b vectors using the NdeI/XhoI sites resulting in plasmids for WT RIα_1-100_, and variants were RIα_1-100_(6R-K) and RIα_1-100_(6R-A), and RIα_1-63_. All plasmids included a TEV-cleavable N-terminus hexa-histidine tag. The sequences of the constructs are given in *Supporting Information*.

### Protein Expression and Purification

Wild type and mutant proteins were expressed in BL21(DE3) in LB media. For isotopic labeling of the protein, a modified M9 minimal media containing 1 g/L ^15^NH_4_Cl and/or 2 g/L ^13^C D-glucose was used. Bacteria were grown in shake flasks at 37 °C to an OD_600_ of 0.8, induced with 0.5 mM IPTG and incubated overnight (∼16 hours) at 18 °C. The cells were harvested by centrifugation at 6000 x g for 20 min, resuspended in lysis buffer (20 mM Tris, 300 mM NaCl, 0.01% NaN_3_, pH 7) at 25 °C, lysed by sonication and the cell lysates were cleared by centrifugation (1 hr, 20,000 x g). The cleared supernatant was loaded on gravity-flow column containing 3 mL Ni-sepharose 6 Fast Flow resin (GE Healthcare), washed with 10 column volumes of lysis buffer + 10 mM imidazole and eluted with lysis buffer + 300 mM imidazole. The eluted protein was digested with TEV protease (1% m/m, TEV/total protein), while dialyzed against lysis buffer without imidazole. The digest was applied to the same Ni-sepharose column as previously, and the flow-through was collected. The protein was concentrated to a concentration of ∼15 mg/mL and further purified by size exclusion chromatography using a Superdex 200 Increase 10/300 GL column (GE healthcare) equilibrated in 20 mM HEPES (pH 7), 150 mM KCl, 0.01% NaN_3_. The final purity of the sample was ascertained by running the samples on an SDS–PAGE. U-^13^C,^15^N-labeled RIα_1-100_ was further dialyzed to 50mM sodium acetate (pH 4), 150 mM NaCl to acquire the triple resonance experiment data.

### Preparations of smAKAP for NMR titrations

The peptide smAKAP (TVILEYAHRLSQDILCDALQQWAC) was obtained from Genscript Inc. at 95% purity. The peptide was dissolved in DMSO, followed by 5 min. sonication in a water bath, and stored at −20 °C until use. The peptide stock diluted in 20mM HEPES (pH 7), 150mM KCl, 0.01% NaN_3_ for NMR experiments.

### NMR data collection and processing

For uniformly ^13^C and/or ^15^N labeled RIα_1-100_ (residue 1–100) a series of 1D ^1^H, 2D ^1^H-^15^N heteronuclear single quantum coherence (HSQC) and ^1^H-^15^N transverse relaxation optimized spectroscopy (TROSY)-HSQC spectra were recorded at various pH (pH 4, 5, 6, and 7) and temperature (278 K, 288 K, 298 K and 310 K). The assignment experiments for RIα_1-63_ were recorded on a 700 MHz ^1^H Bruker AVANCE III spectrometer with a cryogenic probe head with sample in a shaped tube. All other NMR experiments were recorded on a 950 MHz ^1^H Bruker NMR spectrometer equipped with cryogenic probe head with pulse field gradient capability with sample in 5 mm Shigemi tubes. Spectra at the intermediate pH values (5 and 6) were used for transfer of assignment from pH 4 to pH 7. For NMR assignment experiments, a uniformly ^15^N and ^13^C labeled RIα_1-100_ was concentrated to 0.4 mM in pH 4 buffer. For the backbone resonance assignment of RIα_1-100_, best-TROSY-HNCACB, best-TROSY-iHNCACB, best-TROSY-HNCO and best-TROSY-HN(CA)CO^43^ were acquired at 310 K. For the backbone resonance assignment of RIα_1-63,_HNCACB,^44^ HN(CA)CO,^45^ and HNCO^46^ were acquired for a sample in 20mM HEPES (pH 7), 150mM KCl, 0.01% NaN_3_, 100 µM DSS at 298 K. All spectra were chemical shift referenced to DSS, processed using NMRPipe and analyzed using NMRFAM-SPARKY and POKY.^47^ The backbone chemical shifts of RIα_1-63_ and RIα_1-100_ were deposited to BMRB with accession number (RIα_1-63_: 26360, RIα_1-100_: 53210). The backbone chemical shifts of RIα_1-63_ were deposited to using pyPECAN^47^ to analyze the secondary structure of RIα_1-63_.

^15^N R relaxation rates were measured using a TROSY-based pulse sequence^48^ at 298 K using 200 µM protein samples prepared in 20mM HEPES (pH 7), 150mM KCl, 0.01% NaN_3_. R relaxation rates were determined using a ^15^N CPMG (Carr-Purcell-Meiboom-Gill) pulse sequence. A series of eleven relaxation delays ranging from 16 ms to 160 ms were employed. Each experiment was done in duplicate to ensure data reproducibility. R relaxation rates were extracted by fitting a mono-exponential decay function to the peak intensities using POKY.

### NMR titration experiments

Uniformly ^15^N-labeled RIα_1-63_ and RIα_1-100_ were titrated with smAKAP, and a 2D ^1^H-^15^N HSQC spectra were recorded at each step. The induced chemical shift perturbations (CSP) at a saturating 1:2 protein:smAKAP molar ratio was calculated using the formula,

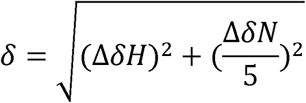

where ΔδH and ΔδN denote the chemical shift changes (in parts per million) in the ^1^H and ^15^N dimensions, respectively.

### In vitro liquid droplet assays

All condensate formation assays were performed in 20 mM HEPES (pH 7.0), 150 mM KCl, 5 mM MgCl_2_,1 mM EGTA, 1 mM DTT, 0.5 mM ATP, and 100 mg/ml polyethylene glycol 4000 (unless specified). Purified proteins were incubated at room temperature for 1 h and imaged using an Olympus IX71 differential interference contrast microscope (DIC) in collagen coated glass bottom dishes (MatTek Corp. Ashland USA). The right-angle light scattering (RALS) and OD_400nm_ RIα_1-63_ and RIα_1-100_ were measured at concentrations of protein and PEG4000 using a *Labbot* instrument (Probation Labs AB, Sweden) at room temperature.

### Fluorescence microscopy and fluorescence recovery after photobleaching (FRAP)

RIα_1-100_was fluorescently labelled by incubation overnight with Atto-488-NHS ester (ATTO-TEC GmbH, Germany) at a 1:2 molar ratio in 20 mM HEPES (pH 8), 150 mM KCl, and 0.01% NaN over night at room temperature. Unreacted dye was removed by dialyzed in 20 mM HEPES (pH 7), 150 mM KCl, and 0.01% NaN□.

Phase separation was induced by adding 100 mg/ml PEG4000 and incubating the sample at 25°C for 1 h. A 30 µL sample was mounted in a MatTek glass-bottom dish (35 mm with a 10 mm #1.5 coverslip microwell) coated with 5% (W/V) polyvinyl alcohol (PVA) to minimize wetting of the droplets. After observing phase separation of RIα_1-100_via fluorescence microscopy, a small droplet region was photobleached with the confocal laser. FRAP experiments were conducted on an inverted Zeiss LSM980 Airyscan 2 confocal microscope. Images were acquired using a 63x oil objective (Plan-Apochromat NA = 1.4) and a 488 nm laser beam at 0.2% laser power at room temperature and detected between 499-552 nm using a PMT detector measuring with a gain of 750 V and an offset of 0%.

For photobleaching, the 488 nm laser line was offset to 100% to bleach a small region of interest (ROI) within phase-separated protein droplets. Pre- and post-bleach images were acquired using the same laser settings at 0.5-second intervals for 2 minutes. The fluorescence recovery of the bleached region was monitored at a resolution of 512 × 512 pixels, 16-bit depth, with a final pixel size of 0.043 µm/pixel. Fluorescence recovery was measured over time, and recovery curves were generated from the average fluorescence intensity within the ROI. Recovery half-times (t_1/2_) were determined by fitting the data to a single exponential recovery model.

### Hydrodynamic Radius measurements by DLS and FIDA

Hydrodynamic radii of RIα_1-63_ and RIα_1-100_ were measured using Dynamic Light Scattering (DLS) with Prometheus Panta (NanoTemper Technologies). Proteins were prepared at concentrations at 50 μM, 100 μM, 200 μM, and 400 μM in 20 mM HEPES (pH 7), 150 mM KCl, and 0.01% NaN□. For each sample, 10 consecutive DLS measurements were conducted at 25 °C using 100% laser power.

Hydrodynamic radii of Atto488-labelled RIα_1-63_ (100 nM) and RIα_1-100_ (250 nM) were measured in the same buffer as DLS using flow induced dispersion analysis (FIDA) using a *Fida 1* instrument (Fida Biosystems ApS, Copenhagen, Denmark). FIDA measurements used PEG-coated capillaries (inner diameter 75 µm, outer diameter 375 µm, total length 100 cm, detection window 84 cm). The capillary was equilibrated with buffer at 1500 mbar for 300 seconds. Analyte solution (buffer) was then introduced at 1500 mbar for 45 seconds, followed by injection of Atto488-tagged RIα_1-63_ and RIα_1-100_ at 50 mbar for 10 seconds. The subsequent analysis was run at 400 mbar for 180 seconds. Taylorgrams were analyzed to determine the apparent hydrodynamic radius of the Atto488-tagged RIα_1-63_ and RIα_1-100_ using the FIDA software suite (version 1.1).

### Atomistic simulations of single dimers

Atomistic simulations were performed using Gromacs 2024.1^49^ and the CHARMM36M force field with scaled TIP3P water for improved treatment of disordered domains in comparison to CHARMM36^50,51^. For all simulations, the time step was 2 fs. Where pressure coupling was used, the pressure was set to 1.0 bar with a compressibility of 4.5×10^−5^ bar^−1^. During equilibration the C-rescale barostat was used, and production simulations were performed using the Parrinello−Rahman barostat with isotropic pressure coupling.^52,53^ Simulation temperature was controlled with the v-rescale thermostat.^54^ Electrostatic interactions were treated with PME with a cut off of 1.2 nm.

Initial structures of a single dimer of RIα_1-100_ were generated using AlphaFold 2 multimer^55^. As these structures showed overestimated helicity in the known disordered regions, they were first subjected to a high temperature unfolding simulation at 450 K for 1 μs, with position restraints on the backbone atoms of the known folded region (residues 12-60). This simulation followed minimization and equilibration in NVT and NPT ensembles. From the high temperature unfolding simulation, single frame starting structures were extracted from the last 400 ns of the trajectory for simulations at 298 K. Starting structures were placed in the centers of dodecahedral simulation boxes separated by 6 nm from the edges. The systems were solvated with 0.15 M NaCl, energy minimized, subject to NVT and NPT equilibration for 1 ns each, and subsequently production run simulations were run for 1 μs. Four replicates were performed for a total simulation time of 4 μs. During equilibration simulations, position restraints were applied on heavy backbone atoms of the protein.

### Coarse grained simulations of single dimers using Martini

From atomistic simulations, the final frame of a production run was used to generate the CG Martini 3 topology of RIα_1-100_ using Martinize2 version 0.11.^56,57^ The latest version of the Martini Go model was used to generate secondary and tertiary structure bonds in the folded regions of the dimer. The Martini Go model describes specialized parameters for the disordered regions, also specified using Martinize2, principally increasing the backbone-water interaction in disordered regions by 0.5 kJ/mol.^58^ In the central folded domain, the backbone-water interaction was not modified. Appropriate side chain fixes were applied to the structure, again determined by the disordered region input to Martinize2.^59^ Once the CG structure and parameters were generated, single dimer simulations were carried out similarly to atomistic simulations. Structures were solvated and charge-neutralized, with ions added to a concentration of 0.15 M NaCl, and energy minimized, equilibrated. Subsequently a production run was simulated for 10 μs. Three repeats of this simulation were performed.

Simulations used the standard Martini 3 input parameters. Electrostatics were treated with reaction field with a cutoff of 1.1 nm, and van der Waals interactions were treated with potential-shift-Verlet, also with a cutoff of 1.1 nm^60^ and the recommended settings to correct neighbourlist artifacts.^61^ Pressure was maintained isotropically at 1 bar using the C-rescale barostat during equilibration and the Parrinello−Rahman barostat for production, respectively. Compressibility was set to 3.0×10^−4^ bar^−1^. The v-rescale thermostat was used to maintain temperature at 300 K for all simulations. During equilibration, position restraints were applied to the backbone of the protein.

### Coarse-grained simulations of condensates using Martini

Initial configurations of 40 RIα_1-100_ dimers were generated using Gromacs version 2024.1 from sampling the concatenated trajectories of the single dimers. Initially, dimers were placed equally spaced in a rectangular box in a 2×2×10 arrangement, solvated and the salt concentration was set to 0.15 M NaCl in addition to neutralizing ions. The systems were subsequently energy minimized and equilibrated. After equilibration, production simulations were performed for 20 μs. Simulation input parameters were identical to single chain simulations, with the exception of the use of semi-isotropic pressure coupling (compressibility = 0 in the xy plane) for equilibration and production runs. Three replicates were constructed and simulated in this manner. In an additional setup, the 40 dimers were pre-assembled into a slab geometry, followed by the same equilibration and production steps.

### Simulation analysis

Analysis of simulations was performed using custom scripts in Python and the MDAnalysis package^62,63^. To comparably calculate contacts, atomistic trajectories were mapped to Martini resolution using fast forward.^64^ Contacts between residues were defined as two backbone beads of proteins at distances within 6 Å. Visualization of MD simulations was done with VMD,^65^ and Martini simulations made use of the MartiniGlass package in Python.^66^

## RESULTS

### The interdomain linker is crucial to PKA RIα phase separation

To identify the molecular determinants of the minimal phase-separating component within RIα, we examined the phase-separating properties of two RIα protein constructs (see Fig. 1A): The first (RIα_1-100_) included the N-terminal D/D domain and the linker region, while the other (RIα_1-63_) consisted only of the D/D domain. Residues 62–100 form an intrinsically disordered region (IDR), which includes an inhibitory sequence that folds upon binding to the catalytic subunit^9^. Neither construct showed signs of phase separation at concentrations up to 300 µM with physiological salt concentration, although a similar construct forms condensates in HEK293T cells.^16^ To reveal latent propensity for phase separation, we also performed experiments in the presence of a crowder (PEG4000). In the presence of 50 mg/ml PEG4000, Atto488-tagged RIα_1-100_ formed spherical protein droplets measuring 3–5 µm in diameter, whereas RIα_1-63_ remained soluble (Fig. 1B).

To explore their phase behavior, RIα_1-63_ and RIα_1-100_ were investigated at varying protein and PEG4000 concentrations using static light scattering. For 25 µM RIα_1-100_, scattering was noticeably increased even at the lowest crowder concentration tested (50 mg/ml) and further increased with the amount of crowder (Fig. 1C) and RIα_1-100_ (Fig. 1D). In contrast, scattering from samples of RIα_1-63_ remained consistently low, and we did not see turbidity at any combination of protein or crowder concentration (Fig. 1C,D and Fig. S1A,B).

The temperature dependence of light scattering for RIα_1-100_ revealed an initial decrease in scattering followed by a sharp increase at ∼60°C (Fig. S1C), which was irreversible upon cooling. We attribute this behavior to dissolution of condensates around room temperature followed by aggregation of the protein at high temperature due to unfolding of the D/D domain. Such traces are not suitable for quantitative analysis but indicate *upper critical solution temperature* (UCST) phase behavior. We investigated the fluidity of RIα_1-100_ droplets using FRAP, which shows ∼80% recovery with a t_½_ of ∼5 s indicating a high mobility of proteins in the droplet (Fig. 1E). Collectively, the results indicate that RIα_1-100_phase separates into dynamic condensates at low concentrations of crowder, and that this behavior depends decisively on the presence of the IDR.

### NMR shows interactions between the IDR and the D/D domain

We used NMR spectroscopy to characterize RIα_1-100_ and RIα_1-63_ that do and do not phase separate, respectively. We recorded TROSY-based ^1^H-^15^N 2D HSQC spectra of U-^13^C/^15^N-labeled RIα_1-100_ at 310 K and pH 4, 5, 6 and 7 (Fig. S2A). We found that, at pH 4, RIα_1-100_ displayed a subset of weak but well-dispersed peaks alongside strong cross peaks that clustered in a narrow region around 8 ppm. Such observations are explained by well-distributed signals from the folded D/D domain, superposed by sharp overlapped peaks from the disordered region. At higher pH values, many of the D/D domain peaks broadened and disappeared (Fig. S2A). In marked contrast, RIα_1-63_ displayed well-resolved peaks with uniform intensity at all pH values (Fig. S3), expected for a small folded, globular proteins in agreement with earlier studies of bovine RIα_12-61_.^67^ These results forebode that the longer construct must harbor slow internal dynamic interactions that lead to line broadening (*vide infra*).

We assigned the NMR resonances using triple resonance experiments and obtained assignments for 57 of 61 expected NH correlations in the 2D ^1^H^−15^N HSQC spectrum of RIα_1-63_ (Fig. S3A). 55 out of 61 assignments could then be successfully transferred to the ^1^H^−15^N HSQC spectra of the longer construct RIα_1-100_ (Fig. 2A). Inference of the assignments in this manner was imperative, as the spectral quality for the folded domain in the longer context was poor due to dynamic effects that resulted in many missing correlations in the 3D spectra. The IDR has sufficiently intense peaks to allow its direct NMR assignment at pH 7, and 27 correlations were observed for the IDR using standard 3D triple resonance experiments. In this manner, 82 out of 93 expected NH resonances could be assigned for RIα_1-100_ at pH 7, in spite of its severely broadened spectrum (Fig. S2D).

**Figure 2:**
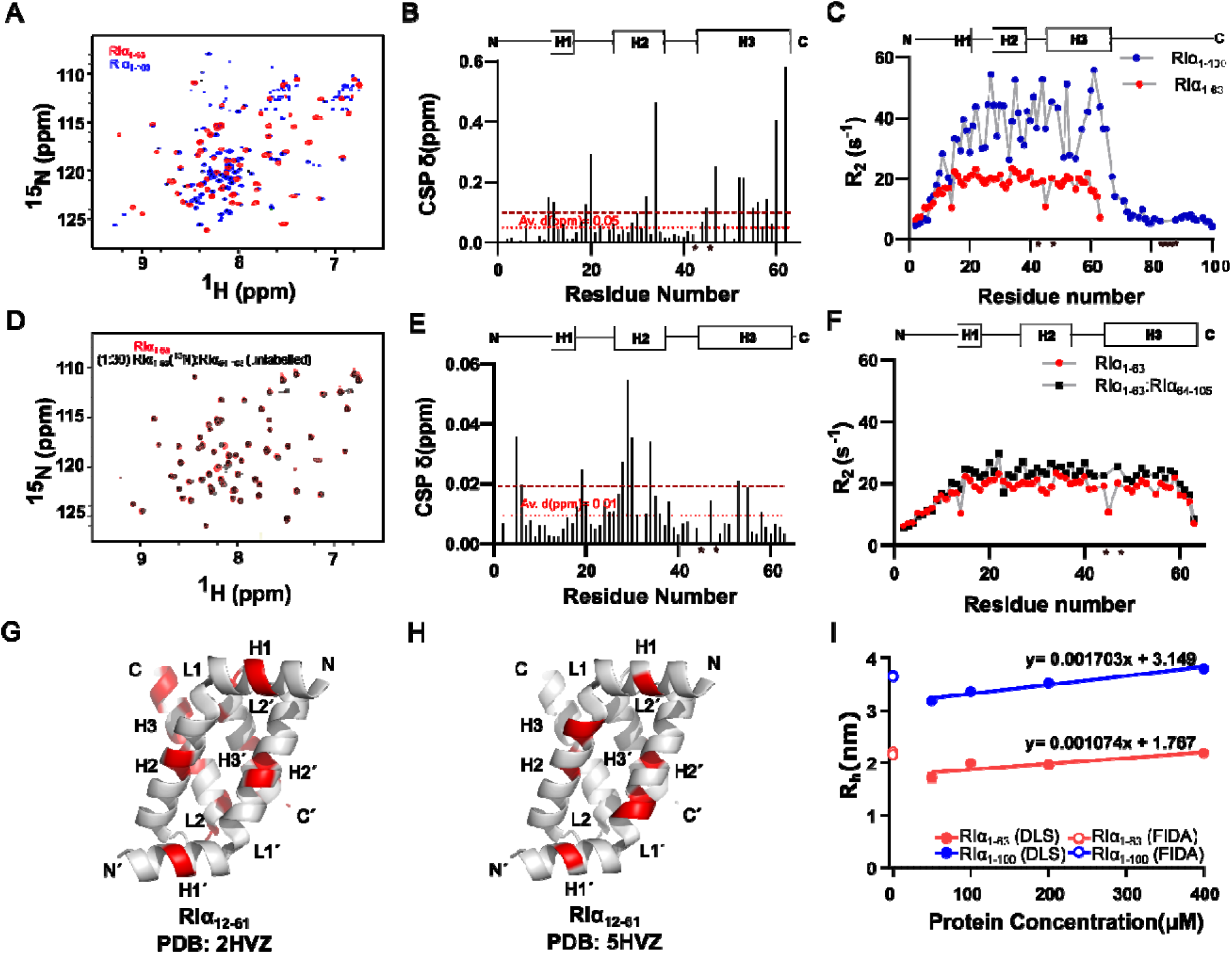
IDR tail interaction with folded D/D domain. A) Overlaid 2D ^1^H-^15^N HSQC spectra of RIα_1-63_ (red) an RIα_1-100_ (blue). B) CSP plot for RIα_1-63_ compared with RIα_1-100_. The red line in-dot represents the average CSP. Th red line in-dash represents two times the average CSP. C) R_2_ plot of RIα_1-63_ (red) and RIα_1-100_ (blue) D) Overlaid 2D ^1^H-^15^N HSQC spectra of free RIα_1-63_ (red) and RIα_1-63_-RIα_64-105_ (black) complex at a 1:30 protein to peptide molar ratio. E) CSP plot of RIα_1-63_-RIα_64-105_ complex at a 1:30 protein to peptide molar ratio. The red line in-dot represents the average CSP. The red line in-dash represents two times the average CSP. F) R_2_ plot of free RIα_1-63_(red) and RIα_1-63_ - RIα_64-105_ complex (black). Interactions are mapped on the RIα_12-61_ structure (PDB: 5hvz^70^). Data above two times the average CSP are shown in red. G) Residues with significant CSPs when including the IDR in the construct (panel B) are mapped on the RIα_12-61_ structure. H) Residues with significant CSPs upon adding the tail IDR *in trans* (panel E) are mapped on the RIα_12-61_ structure. I) Hydrodynamic radius of RIα_1-63_ and RIα_1-100_ measured by FIDA and DLS.

Differences in structure between the two constructs were first investigated by a comparison of backbone chemical shifts under identical conditions. The largest chemical shift perturbations (CSPs) were found in a junction between the IDR and the D/D domains (Figure 2B). These differences are explained by the presence of a charged carboxy terminus in the shorter construct that is not present in the long construct. However, additional perturbations were observed throughout the protein, which must arise from an interaction between the IDR and the D/D domain. To test this, we conducted backbone ^15^N NMR relaxation experiments that compare the dynamic properties of RIα_1-63_and RIα_1-100_. As expected, RIα_1-63_showed uniform transverse relaxation rates (R_2_), consistent with a folded globular domain (Fig. 2C). In contrast, RIα_1-100_ exhibited two regions with distinct R_2_ values (Fig. 2C): R_2_ rates were uniformly low for the linker region, which is typical for disordered regions. However, for the D/D domain, the ^15^N R_2_ values are much higher than for the corresponding residues in RIα_1-63_ suggesting either exchange broadening resulting from transient interactions on a microsecond to millisecond time scale or the formation of larger complexes comprising of multiple protein copies.

To probe the presence of a potential interaction between the D/D domain and the IDR, we made a peptide corresponding to the C-terminal IDR alone (RIα_64-105_) and titrated it into ^15^N-labeled RIα_1-63_. The linker does not have any aromatic residues, so the construct was extended to encompass Y105 to allow quantification by UV absorption. We observed a gradual shift in the positions of several HSQC cross peaks as the concentration of RIα_64-105_ was increased. This indicates that the interaction between RIα_1-63_ and RIα_64-105_ is fast on the NMR timescale with a bound-state lifetime of microseconds or shorter (Figure 2D). Mapping of the chemical shift perturbations on the sequence revealed a similar pattern of affected residues as in RIα_1-100_, suggesting that a similar interaction takes place *in cis* and *in trans* (Fig. 2E). A comparison of the ^15^N R_2_ rates of the RIα_1-63_:RIα_64-105_ complex with those of free RIα_1-63_showed only a slight increase, indicating weaker interactions when the regions are not tethered; this can be the result of faster dynamics and/or an interaction where the equilibrium lies more towards the unbound form (Fig. 2F).

NMR CSP analysis at a 1:30 ratio of RIα_1-63_to RIα_64-105_ identified key residues in the interaction between the D/D domain and the IDR. Several residues from helix H2 and from the N-terminus showed CSPs more than two times the average, suggesting that a set of residues in the D/D domain are involved in the interaction (Fig. 2E). We mapped NMR CSPs onto the D/D domain homodimer structure (PDB: 5HVZ) to identify interacting surfaces (Fig. 2G,H). CSP analysis comparing RIα variants with and without the IDR (Fig. 2B,G) with the titration using the IDR *in trans* (Figure 2E,H) mainly differ in the involvement of helix H3. Helix 3 is the most C-terminal one and the attachment site of the IDR, and NMR chemical shifts would already be expected to be perturbed in the absence of an interaction. The two CSP mappings agree on an involvement of helices H1 and H2 on the opposite side of the domain to H3.

To test for multimerization as a contribution to the CSPs and elevated ^15^N R_2_ values in R1α_1-100_ compared to RIα_1-63_, we measured the hydrodynamic radius of both constructs using dynamic light scattering (DLS) and flow-induced dispersion analysis (FIDA). The combination of the two techniques allowed us to span a ∼1000-fold difference in protein concentration to probe for potential transient intermolecular interactions: FIDA uses nanomolar protein concentrations, whereas DLS works best at the high micromolar concentrations where the NMR experiments were also conducted. FIDA measurements resulted in hydrodynamic radii (R_h_) of 2.17 ± 0.019 nm for RIα_1-63_ and 3.65 ± 0.010 nm for RIα_1-100_, respectively, while DLS measurements yielded R_h_ values of 1.99 ± 0.04 nm for RIα_1-63_ and 3.36 ± 0.02 nm for RIα_1-100_. The slightly larger R_h_ observed by FIDA can possibly be attributed to the presence of an Atto488-fluorophore in this construct. The similar R_h_ across a 1.000-fold concentration difference suggests that multimerization is not prevalent at the concentrations used for NMR spectroscopy. Furthermore, we analyzed the concentration-dependence of the hydrodynamic radius using DLS measurement to test interparticle interactions. For both RIα_1-63_ and RIα_1-100_, the radius of hydration increases slightly with concentration suggesting overall attractive interparticle interactions. The slope was steeper for RIα_1-100_ (0.0017) than for RIα_1-63_ (0.0011), indicating stronger attraction, in agreement with a stronger tendency to form homotypic phase separation.

Taken together, these measurements suggest that both the IDR and the D/D domain have potential for self-attraction, but that these attractions are not sufficient to lead to formation of multimers at the concentrations used for NMR and DLS.

### The arginine residues within the IS contribute to RIα_1-100_ phase separation

Next, we set out to map the sequence features of the IDR that drive phase separation. The IDR includes the so-called inhibitory sequence (IS), which is disordered in the catalytically active form of PKA and binds the PKA_cat_ subunit to form the inactive PKA holoenzyme. The IS contains four arginine residues (residues 94-97) (Fig. 3A), which are strong drivers of IDR phase separation.^68^ To test whether the arginine residues also drive phase separation of RIα_1-100_, we made three variants: (a) RIα_1-93_ where the IS is absent; (b) RIα_1-100_(6R-K) where the 6 arginine residues in the IDR are changed to lysine, preserving the positive charge but removing the guanidinium group; (c) RIα_1-100_ (6R-A) where the six arginine residues are mutated to alanine (Fig. 3A).

**Figure 3:**
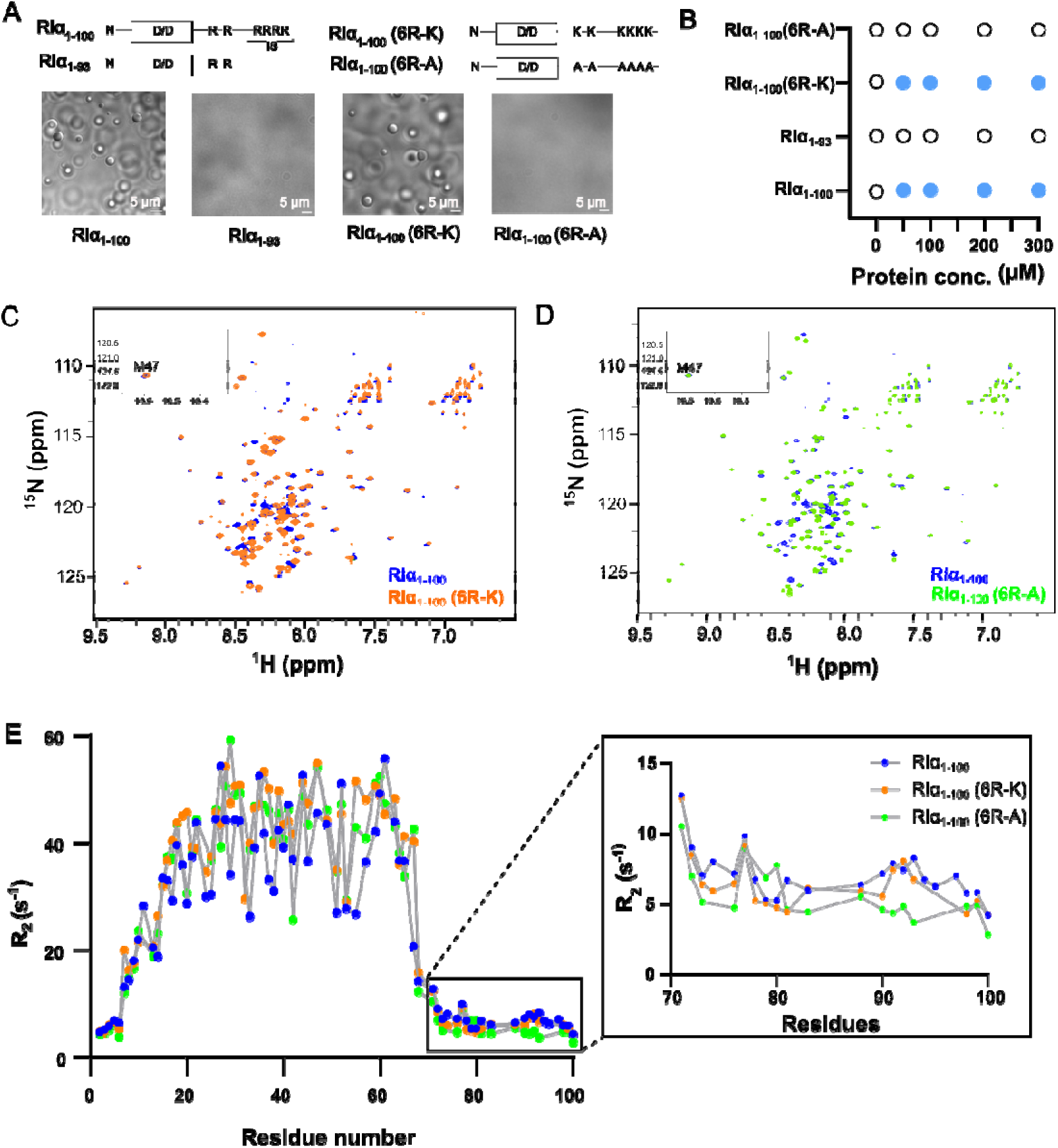
Positively charged residues of the RIα IDR tail drive condensate formation. A) WT type and mutants constructs of PKA R subunit. DIC microscope images of liquid droplets by 100 µM RIα_1-100_, no droplet formation b 100 µM RIα_1-93_ mutant, liquid droplets by 100 µM RIα_1-100_ (6R-K) and no droplet formation by RIα_1-100_ (6R-A) mutant. B) Representative *in vitro* phase diagram of WT type and mutants in presence of 100 mg/ml PEG400 crowder. C) Overlaid 2D ^1^H-^15^N HSQC spectra of free RIα_1-100_ (blue) and RIα_1-100_ (6R-K) (orange) at pH 7, 298 K. D) Overlaid 2D ^1^H-^15^N HSQC spectra of free RIα_1-100_ (blue) and RIα_1-100_ (6R-A) (green) at pH 7, 298 K. E) Overlaid R_2_ plot of RIα_1-100_ (blue), RIα_1-100_ (6R-K)(orange) and RIα_1-100_ (6R-A) (green). Enlarged view shows the R_2_ of the residues (71-100) of IDR tail of RIα_1-100_ (blue), RIα_1-100_ (6R-K) (orange) and RIα_1-100_ (6R-A) (green).

We tested the variants in the presence of an external crowder (PEG4000), where RIα_1-100_ robustly formed spherical condensates (Fig. 3A,B), whereas RIα_1-93_ and RIα_1-100_ (6R-A) failed to form droplets at the concentrations tested (Fig. 3B), nor formed punctae in the presence of much higher concentrations of crowder (Fig. S4A,B). In contrast, RIα_1-100_ (6R-K) formed liquid droplets similar to those observed for RIα_1-100_under these conditions (Fig. 3A,B and S4C). This suggests that the phase separation propensity of the IDR is driven by the arginine residues in the IS.

Previous NMR studies of phase-separating proteins have shown broadening of signals under phase-separating conditions.^21^ Several studies have also shown that proteins with propensity for phase separation form soluble “nanoclusters” even under conditions where macroscopic phase separation is not observed.^69^ The elevated transverse ^15^N relaxation rates observed for RIα_1-100_compared with RIα_1-63_ might be linked to phase-separation propensity. To test this possibility, we recorded 2D ^15^N-^1^H HSQC spectra of RIα_1-100_ (6R-K) and RIα_1-100_ (6R-A) (Fig. 3C, D). New peaks were observed corresponding to lysine and alanine residues respectively, and the arginine correlations were absent. However, the overall appearance of the spectra was similar to those for wild-type RIα_1-100_, with a large variation in peak intensities. To probe the dynamics more quantitatively, we recorded ^15^N R_2_ for both mutant versions. Relaxation rates were similar to wild-type RIα_1-100_ within the D/D-domain, indicating similar dynamics for the folded domain (Fig. 3E). However, the rates for the IDR region showed a noticeable decrease for RIα_1-100_ (6R-A), particularly for residues 70–100, when compared to wild-type and RIα_1-100_ (6R-K). This reduction shows a clear loss of dynamic interactions of the IDR with the D/D domain.

### AKAP binding to the D/D domain competes with the IDR interaction

The D/D domain binds amphipathic helices from A-kinase anchoring proteins (AKAPs) to regulate the spatiotemporal activity of the kinase. AKAPs bind to a hydrophobic patch across helices H1 and H2,^70^ which overlaps with the surface where we observed chemical shift perturbations in the presence of the IDR. This suggests a potential competition between the IDR and smAKAP for binding to the D/D domain. To explore this, we titrated ^15^N-labeled RIα_1-63_ and RIα_1-100_ with increasing concentrations of smAKAP and monitored the interaction using 2D ^1^H-^15^N HSQC NMR. The interaction of RIα_1-63_ with smAKAP is in a slow exchange on the NMR time scale, with the disappearance of free protein signals and concomitant appearance of new peaks. At 1:2 molar ratio RIα_1-63_:smAKAP, no free protein peaks remained (Fig. 4A), in line with a strong interaction that is expected (*K*_d_ = 7 nM).^70^ For RIα_1-100_, most of the well-dispersed amide peaks disappeared, but the peaks for the disordered residues remained unaffected (Fig. 4B). Whereas the NMR signals of the D/D were lost, the dynamic properties of the IDRs can still be analyzed in the context of bound smAKAP. Chemical shift perturbations suggest changes throughout the D/D domain in RIα_1-63_ upon smAKAP binding. The few peaks that are retained in this region in RIα_1-100_ have roughly similar CSPs (Fig. 4C). Additionally, the IDR is also affected by the binding of smAKAP as demonstrated by small, but significant values in this region. ^15^N R_2_ for RIα_1-100_ revealed reduced rates for residues 70-100 in the complex (Fig. 4D). The decrease in R_2_ rates cannot be explained by a direct interaction between the IDR and the smAKAP, but, instead, suggest a release of interactions that pre-exist between the IDR and the D/D domain.

**Figure 4:**
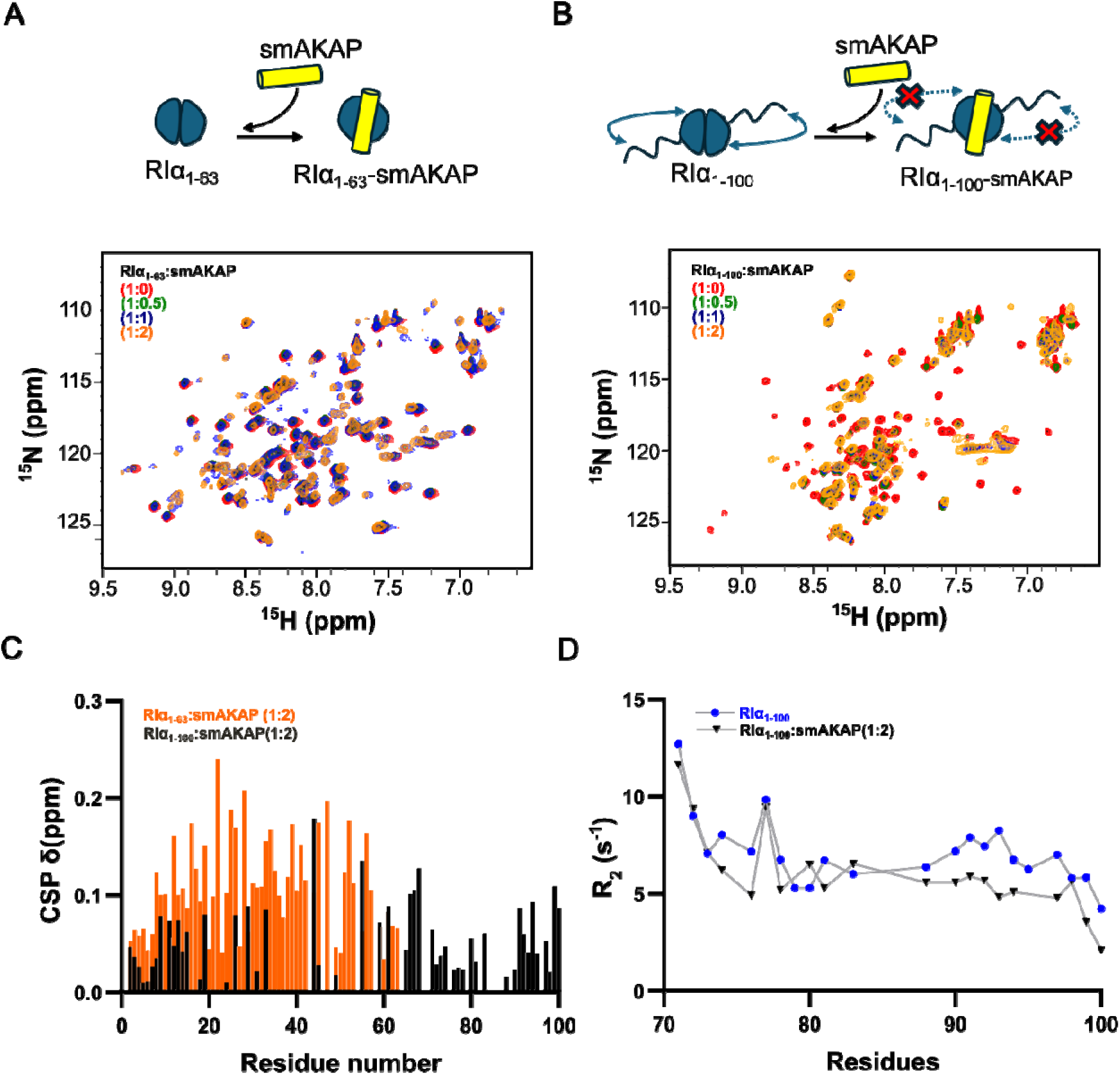
The IDR tail modulates the interaction between smAKAP and the RIα D/D domain. A) Schematic representation of smAKAP interaction with RIα_1-63_. Overlaid 2D ^1^H-^15^N HSQC spectra of free RIα_1-63_(red) and RIα_1-63_-smAKAP (green) complex at a 1:0.5 protein to peptide molar ratio, RIα_1-63_-smAKAP (blue) complex at a 1:1 protein to peptide molar ratio and RIα_1-63_-smAKAP(orange) complex at a saturated 1:2 protein to peptide molar ratio. B) Schematic representation of smAKAP interaction with RIα_1-100_ which contains the IDR tail. Overlaid 2D ^1^H-^15^N HSQC spectra of free RIα_1-100_ (red) and RIα_1-100_-smAKAP (green) complex at a 1:0.5 protein to peptide molar ratio, RIα_1-100_-smAKAP (blue) complex at a 1:1 protein to peptide molar ratio and RIα_1-100_-smAKAP (orange) complex at a saturated 1:2 protein to peptide molar ratio. C) CSP plot of RIα_1-63_-smAKAP complex (orange) and RIα_1-100_-smAKAP complex (black) at the saturated 1:2 protein to peptide molar ratio. D) Overlaid R_2_ plot of residues (71-100) of IDR tail of free RIα_1-100_ (blue) and RIα_1-100_-smAKAP complex (black) at the saturated 1: protein to peptide molar ratio.

### Multiscale simulations of RIα_1-100_ dimers reveal extensive interdomain interactions

To shed further light on the intra- and intermolecular interactions revealed by the NMR studies, we performed MD simulations of RIα_1-100_, using a multiscale approach that combines atomistic and CG simulations. Initially, we compared simulations of RIα_1-100_ dimers performed using the atomistic CHARMM force field with CG simulations using the Martini3 force field. Across both resolutions, the global dimensions of the PKA RIα_1-100_ were similar, with the radius of gyration of the atomistic model 2.30 +/− 0.19 nm, comparable to 2.58 +/− 0.03 nm for the CG one. Furthermore, the intramolecular contact maps showed similar contact patterns and prevalences (Fig. 5A), conveying that the CG simulations reproduce the conformational ensemble of the protein well. The contact maps show pervasive self-interaction within the IDR and between the IDR and the D/D domains, corroborating the experimental results. Within each monomer, the most prevalent contacts are between the disordered domains and the N-terminus of the folded domain (Fig. 5B).

**Figure 5:**
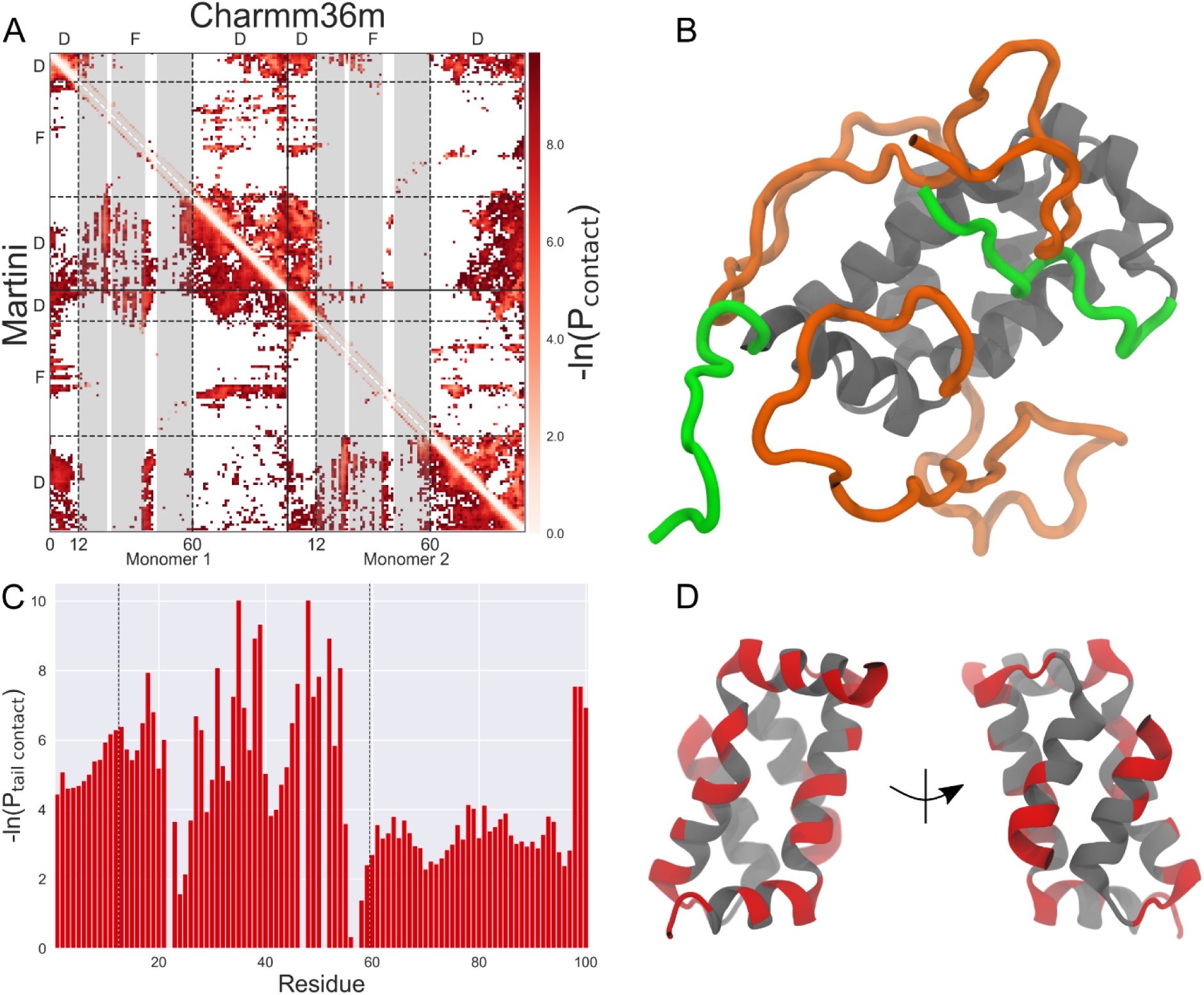
Multiscale simulations of single RIα dimers. A) Contact maps of simulations at atomistic (upper triangle) and Martini (lower triangle) resolution. The dashed lines indicate the folded (F) and disordered (D) regions of both monomers. Atomistic trajectories have been mapped to Martini resolution for comparability. B) Snapshot of the PKA RI dimer during the atomistic simulation, viewing the N-terminal face with contacts from the disordered C-terminus. The N-terminal disordered region is colored in green and the C-terminus disordered region in orange. The central folded domain is shown in grey. C) Contacts between residues in the disordered C terminus and all other residues for a single monomer at the CG resolution averaged over the whole molecule. Dashed lines indicate the disordered/folded domain boundaries. D) Prevalent contacts found in C) in the folded domain (residues 12 to 60). Left, N-terminal face, right, C-terminus face.

The interactions between the IDR and the folded domain were mapped back onto a single monomer (Fig. 5C). As with the experimental results, the solvent-exposed residues in the N-terminal helices appear to all have significant interactions with the disordered tail, along with the short interhelical loop between residues 40 and 44. The contacts found between the disordered and folded domains largely agreed with those found by NMR (Fig. 2G,H). In particular, the simulation showed prevailing contacts between the disordered region and solvent-exposed residues in helix H2 of the monomers. Further, these contacts were most strongly associated with residues Asp33 and Glu55 of this helix, suggesting that favorable electrostatic interactions play an especially important role in the cross-domain interaction.

### Coarse-grained MD simulations reveal disordered-folded interactions significant for phase separation

The Martini3 force field was used to simulate the spontaneous phase separation of 40 RIα_1-100_dimers in triplicate. The initial configurations of RIα_1-100_ dimers were generated from the single dimer simulations described above (Fig. 6A). After a 20 µs production simulation, all three repeats converged to a largely phase-separated state (Fig. 6B) where the majority of dimers were in one assembly and the rest in a smaller assembly.

**Figure 6:**
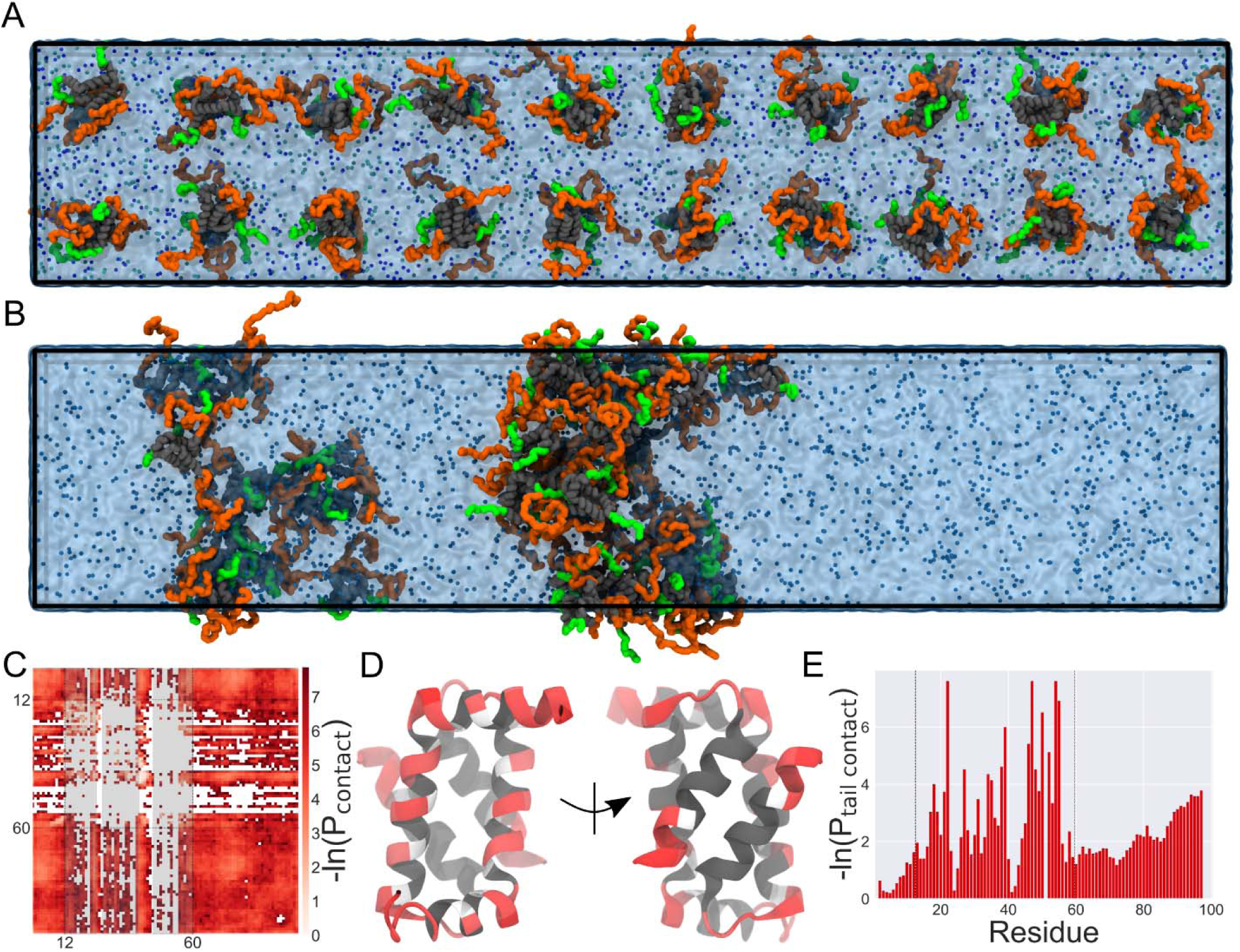
Simulations of self-associating RIα condensates. Condensate simulations are set up from equally space dimers (A) and run in production for 20 μs (B). After 20 μs, the dimers have formed two phase-separated droplets. The N/C-terminal disordered regions are colored in green/orange, the central folded domain is shown in grey. Ions are depicted as blue spheres, and water as a transparent blue surface. C) Contacts formed between dimers across the last 4 μs of the simulation. Contacts here are defined as inter-dimer contacts and mapped back onto a single monomer. D) Contacts between the disordered C termini and the folded domains of other dimers in the condensate. E) Contacts between dimer units averaged over the whole molecules.

Similar to the single-molecule contacts characterized above, we calculated inter-dimer contacts across the final 4 μs of simulation time – i.e. excluding inter-protomer contacts within the same dimer. The contacts (Fig. 6C) are dominated by disordered-disordered contacts with little apparent residue specificity. However, there are also a number of disordered-folded contacts between molecules following the same patterns as observed for the intra-dimer contacts, where the IDR mainly interacts with exposed residues of the N terminus of helix 2 (Fig. 6D). Again, there is prevalence for charged residues in these interactions, confirming that the arginine residues in the IS play an important role in driving the phase separation of the dimer. Less prominent are folded-folded contacts, which are mostly focused on the charges in the interhelical loop (residues 40-44). To determine the effect that the lack of convergence of the system into a single droplet had on the analysis, we performed further simulations of dimers pre-assembled into slab condensates (Fig. S5); this resulted in identical contacts.

Overall, the contact analysis suggests that phase separation of RIα_1-100_ is driven by intermolecular interactions of the IDR, which engages with both the D/D domain and IDR of other RIα dimers.

## DISCUSSION

Here, we investigated the structure and dynamics of the minimal phase-separating unit of PKA RIα by NMR and MD simulations. Our initial NMR studies of RIα were hindered by surprisingly poor spectral properties. The dimeric D/D domain has a molecular weight of ∼12 kDa and would be expected to tumble rapidly and give sharp and resolved NMR signals. The many weak and broad signals suggested that pervasive conformational exchange on the µs-ms time scale might be present, something that was not observed in previous studies of the D/D domain alone.^67^ Since such broadening can be either intra- or intermolecular in origin, we investigated this by making various constructs and mutants in which regions were deleted or altered, and studying them by NMR spectroscopy and hydrodynamic methods. It was established that the folded D/D domain did not display the broadening and the addition of the subsequent IDR was responsible for the effect. Addition of the IDR *in cis* or *in trans* both revealed the interaction and hydrodynamic measurements excluded intermolecular interactions (dimerization) as the origin. Furthermore, mutations in the IDR that were shown to disrupt phase separation did not improve the NMR properties for the D/D domain, albeit that a slight decrease of transverse relaxation rates was seen for the IDR.

We then turned our attention to the functional role of the IDR. Intrinsically disordered linkers are rarely just passive tethers but also regulate the interactions of the folded domains they flank.^71^ The flanking regions in linkers may thus be presented at effective concentrations in millimolar range,^72^ and even weak interactions are brought into play. Flanking IDRs can either increase affinity, when the IDR interacts favorably with flanking regions in a binding partner, or decrease the affinity by competing with the binding partner for the interface. Such interactions can modulate affinity by several orders of magnitude.^73^ For RIα, the IDR following the inhibitory segment (residues 100-120) was previously found to interact with the cyclic nucleotide binding domain to allosterically regulate the equilibrium between binding competent and incompetent conformations.^20^ Here, we now show that the IDR preceding the IS and the IS itself contact the AKAP binding surface of the D/D domain. In the absence of other interactions, such competing interactions will act to lower the affinity for AKAPs. At the same time, as the interaction is weak, promiscuous interactions with flanking regions in the AKAP may counteract this effect.

Finally, we addressed the question of how the IS and IDR may be involved in the regulation of PKA function at the molecular level; Interactions that drive phase separation can often be observed by NMR studies of the dilute phase, for example the case of TDP-43, where the key interaction hotspots were identified as elevated R_2_ values and chemical shift perturbations.^22,74^ Even when macroscopic phase separation is not observed, “nanoclusters” may still occur^69^ which may be reflected in the NMR spectra.^21^ Moreover, phase separation can be exclusively driven by multivalent interactions between proteins,^75^ where IDRs often contribute with unspecific interactions involving charged and aromatic residues^68^. In a recent study, Hardy et al. introduced mutations into RIα and quantified phase separation in cells.^19^ Focusing on the role of the IDRs, phase separation could be disrupted by removing all charged residues in the linker or by introducing a phosphorylation site in the IS. The mutations used here are complimentary, as they primarily target the arginine residues found in the IS. A model is proposed, where the phase-separation propensity can be disrupted by interfering with the interactions of IDRs and it suggests an important role for the region surrounding the IS. The latter is important as it provides a mechanism for how cAMP increases phase separation propensity.^16,19^ The advantage of working with a minimal construct in vitro is that we can provide a microscopic and quantitative understanding of these interactions by NMR and MD simulations. Both techniques locate the main contact point for the IDR at the AKAP binding face of the D/D domain, with contacts found throughout and involving a large number of residues. The simulations allow us to compare intra- and intermolecular interactions, which are remarkably similar (Fig. 5D, 6D) and suggest that intramolecular interactions compete with similar intermolecular ones. This points to an analogy with homotypic phase separation of IDPs and expands it to encompass ligand-induced competition: Proteins with IDRs can form or dissolve condensates in response to small molecule cues that – at the molecular level – rely on very similar interactions but result in a completely different outcome at the mesoscopic and cellular level.

## Acknowledgements

This work was supported by Novo Nordisk Foundation grant NNF20OC0063808, ‘BOUNDLESS’ to F.A.A.M., M.K. and S.J.M and equipment grants CF20-0610 & CF21-0164 from the Carlsberg Foundation). Access to the NMR spectrometers at the Danish Center for Ultrahigh-Field NMR Spectroscopy (Ministry of Higher Education and Science grant AU- 2010- 612-181) is gratefully acknowledged. We thank SURF (www.surf.nl) for the support in using the National Supercomputer Snellius. We would like to thank Lisbeth Schmidt Laursen, Mette Hoffmann Asmussen, Dennis Wilkens Juhl and Jan S. Nowak for technical support.

## Notes

### Competing Interest Statement

The authors have declared no competing interest.

## References

(1) Kemp, B. E.; Graves, D. J.; Benjamani, E.; Krebs, E. G. Role of Multiple Basic Residues in Determining the Substrate Specificity of Cyclic AMP-Dependent Protein Kinase. J. Biol. Chem. 1977, 252 (4), 4888–4894.

(2) Isobe, K.; Jung, H. J.; Yang, C.-R.; Claxton, J.; Sandoval, P.; Burg, M. B.; Raghuram, V.; Knepper, M. A. Systems-Level Identification of PKA-Dependent Signaling in Epithelial Cells. Proc. Natl. Acad. Sci. 2017, 114 (42). 10.1073/pnas.1709123114.

(3) Boularan, C.; Gales, C. Cardiac cAMP: Production, Hydrolysis, Modulation and Detection. Front. Pharmacol. 2015, 6, 203. 10.3389/fphar.2015.00203.

(4) Kandel, E. R. The Molecular Biology of Memory: cAMP, PKA, CRE, CREB-1, CREB-2, and CPEB. Mol. Brain 2012, 5 (1), 14. 10.1186/1756-6606-5-14.

(5) Jhala, U. S.; Canettieri, G.; Screaton, R. A.; Kulkarni, R. N.; Krajewski, S.; Reed, J.; Walker, J.; Lin, X.; White, M.; Montminy, M. cAMP Promotes Pancreatic Beta-Cell Survival via CREB-Mediated Induction of IRS2. Genes Dev. 2003, 17 (13), 1575–1580. 10.1101/gad.1097103.

(6) Taylor, S. S.; Zhang, P.; Steichen, J. M.; Keshwani, M. M.; Kornev, A. P. PKA: Lessons Learned after Twenty Years. Biochim. Biophys. Acta BBA - Proteins Proteomics 2013, 1834 (7), 1271–1278. 10.1016/j.bbapap.2013.03.007.

(7) Taylor, S. S.; Ilouz, R.; Zhang, P.; Kornev, A. P. Assembly of Allosteric Macromolecular Switches: Lessons from PKA. Nat. Rev. Mol. Cell Biol. 2012, 13 (10), 646–658. 10.1038/nrm3432.

(8) Kim, C.; Cheng, C. Y.; Saldanha, S. A.; Taylor, S. S. PKA-I Holoenzyme Structure Reveals a Mechanism for cAMP-Dependent Activation. Cell 2007, 130 (6), 1032–1043. 10.1016/j.cell.2007.07.018.

(9) Hamuro, Y.; Anand, G. S.; Kim, J. S.; Juliano, C.; Stranz, D. D.; Taylor, S. S.; Woods, V. L. Mapping Intersubunit Interactions of the Regulatory Subunit (RIα) in the Type I Holoenzyme of Protein Kinase A by Amide Hydrogen/Deuterium Exchange Mass Spectrometry (DXMS). J. Mol. Biol. 2004, 340 (5), 1185–1196. 10.1016/j.jmb.2004.05.042.

(10) Canaves, J. M.; Taylor, S. S. Classification and Phylogenetic Analysis of the cAMP-Dependent Protein Kinase Regulatory Subunit Family. J. Mol. Evol. 2002, 54 (1), 17–29. 10.1007/s00239-001-0013-1.

(11) Heller, W. T.; Vigil, D.; Brown, S.; Blumenthal, D. K.; Taylor, S. S.; Trewhella, J. C Subunits Binding to the Protein Kinase A RIα Dimer Induce a Large Conformational Change. J. Biol. Chem. 2004, 279 (18), 19084–19090. 10.1074/jbc.M313405200.

(12) Bock, A.; Annibale, P.; Konrad, C.; Hannawacker, A.; Anton, S. E.; Maiellaro, I.; Zabel, U.; Sivaramakrishnan, S.; Falcke, M.; Lohse, M. J. Optical Mapping of cAMP Signaling at the Nanometer Scale. Cell 2020, 182 (6), 1519–1530.e17. 10.1016/j.cell.2020.07.035.

(13) Bock, A.; Irannejad, R.; Scott, J. D. cAMP Signaling: A Remarkably Regional Affair. Trends Biochem. Sci. 2024, 49 (4), 305–317. 10.1016/j.tibs.2024.01.004.

(14) Depry, C.; Allen, M. D.; Zhang, J. Visualization of PKA Activity in Plasma Membrane Microdomains. Mol BioSyst 2011, 7 (1), 52–58. 10.1039/C0MB00079E.

(15) Smith, F. D.; Reichow, S. L.; Esseltine, J. L.; Shi, D.; Langeberg, L. K.; Scott, J. D.; Gonen, T. Intrinsic Disorder within an AKAP-Protein Kinase A Complex Guides Local Substrate Phosphorylation. eLife 2013, 2, e01319. 10.7554/eLife.01319.

(16) Zhang, J. Z.; Lu, T.-W.; Stolerman, L. M.; Tenner, B.; Yang, J. R.; Zhang, J.-F.; Falcke, M.; Rangamani, P.; Taylor, S. S.; Mehta, S.; Zhang, J. Phase Separation of a PKA Regulatory Subunit Controls cAMP Compartmentation and Oncogenic Signaling. Cell 2020, 182 (6), 1531–1544.e15. 10.1016/j.cell.2020.07.043.

(17) Zhang, J. Z.; Mehta, S.; Zhang, J. Liquid–Liquid Phase Separation: A Principal Organizer of the Cell’s Biochemical Activity Architecture. Trends Pharmacol. Sci. 2021, 42 (10), 845–856. 10.1016/j.tips.2021.07.003.

(18) López-Palacios, T. P.; Andersen, J. L. Kinase Regulation by Liquid–Liquid Phase Separation. Trends Cell Biol. 2023, 33 (8), 649–666. 10.1016/j.tcb.2022.11.009.

(19) Hardy, J. C.; Pool, E. H.; Bruystens, J. G. H.; Zhou, X.; Li, Q.; Zhou, D. R.; Palay, M.; Tan, G.; Chen, L.; Choi, J. L. C.; Lee, H. N.; Strack, S.; Wang, D.; Taylor, S. S.; Mehta, S.; Zhang, J. Molecular Determinants and Signaling Effects of PKA RIα Phase Separation. Mol. Cell 2024, 84 (8), 1570–1584.e7. 10.1016/j.molcel.2024.03.002.

(20) Akimoto, M.; Selvaratnam, R.; McNicholl, E. T.; Verma, G.; Taylor, S. S.; Melacini, G. Signaling through Dynamic Linkers as Revealed by PKA. Proc. Natl. Acad. Sci. 2013, 110 (35), 14231–14236. 10.1073/pnas.1312644110.

(21) Murthy, A. C.; Fawzi, N. L. The (Un)Structural Biology of Biomolecular Liquid-Liquid Phase Separation Using NMR Spectroscopy. J. Biol. Chem. 2020, 295 (8), 2375–2384. 10.1074/jbc.REV119.009847.

(22) Conicella, A. E.; Zerze, G. H.; Mittal, J.; Fawzi, N. L. ALS Mutations Disrupt Phase Separation Mediated by α-Helical Structure in the TDP-43 Low-Complexity C-Terminal Domain. Structure 2016, 24 (9), 1537–1549. 10.1016/j.str.2016.07.007.

(23) Ryan, V. H.; Dignon, G. L.; Zerze, G. H.; Chabata, C. V.; Silva, R.; Conicella, A. E.; Amaya, J.; Burke, K. A.; Mittal, J.; Fawzi, N. L. Mechanistic View of hnRNPA2 Low-Complexity Domain Structure, Interactions, and Phase Separation Altered by Mutation and Arginine Methylation. Mol. Cell 2018, 69 (3), 465–479.e7. 10.1016/j.molcel.2017.12.022.

(24) Kim, T. H.; Tsang, B.; Vernon, R. M.; Sonenberg, N.; Kay, L. E.; Forman-Kay, J. D. Phospho-Dependent Phase Separation of FMRP and CAPRIN1 Recapitulates Regulation of Translation and Deadenylation. Science 2019, 365 (6455), 825–829. 10.1126/science.aax4240.

(25) Kim, T. H.; Payliss, B. J.; Nosella, M. L.; Lee, I. T. W.; Toyama, Y.; Forman-Kay, J. D.; Kay, L. E. Interaction Hot Spots for Phase Separation Revealed by NMR Studies of a CAPRIN1 Condensed Phase. Proc. Natl. Acad. Sci. 2021, 118 (23), e2104897118. 10.1073/pnas.2104897118.

(26) Murthy, A. C.; Dignon, G. L.; Kan, Y.; Zerze, G. H.; Parekh, S. H.; Mittal, J.; Fawzi, N. L. Molecular Interactions Underlying Liquid−liquid Phase Separation of the FUS Low-Complexity Domain. Nat. Struct. Mol. Biol. 2019, 26 (7), 637–648. 10.1038/s41594-019-0250-x.

(27) Ambadipudi, S.; Reddy, J. G.; Biernat, J.; Mandelkow, E.; Zweckstetter, M. Residue-Specific Identification of Phase Separation Hot Spots of Alzheimer’s-Related Protein Tau. Chem. Sci. 2019, 10 (26), 6503–6507. 10.1039/C9SC00531E.

(28) Joseph, J. A.; Reinhardt, A.; Aguirre, A.; Chew, P. Y.; Russell, K. O.; Espinosa, J. R.; Garaizar, A.; Collepardo-Guevara, R. Physics-Driven Coarse-Grained Model for Biomolecular Phase Separation with near-Quantitative Accuracy. Nat. Comput. Sci. 2021, 1 (11), 732–743. 10.1038/s43588-021-00155-3.

(29) Tesei, G.; Lindorff-Larsen, K. Improved Predictions of Phase Behaviour of Intrinsically Disordered Proteins by Tuning the Interaction Range. Open Res. Eur. 2023, 2, 94. 10.12688/openreseurope.14967.2.

(30) Borges-Araújo, L.; Patmanidis, I.; Singh, A. P.; Santos, L. H. S.; Sieradzan, A. K.; Vanni, S.; Czaplewski, C.; Pantano, S.; Shinoda, W.; Monticelli, L.; Liwo, A.; Marrink, S. J.; Souza, P. C. T. Pragmatic Coarse-Graining of Proteins: Models and Applications. J. Chem. Theory Comput. 2023, 19 (20), 7112–7135. 10.1021/acs.jctc.3c00733.

(31) Dignon, G. L.; Best, R. B.; Mittal, J. Biomolecular Phase Separation: From Molecular Driving Forces to Macroscopic Properties. Annu. Rev. Phys. Chem. 2020, 71 (1), 53–75. 10.1146/annurev-physchem-071819-113553.

(32) Dignon, G. L.; Zheng, W.; Best, R. B.; Kim, Y. C.; Mittal, J. Relation between Single-Molecule Properties and Phase Behavior of Intrinsically Disordered Proteins. Proc. Natl. Acad. Sci. 2018, 115 (40), 9929–9934. 10.1073/pnas.1804177115.

(33) Dignon, G. L.; Zheng, W.; Kim, Y. C.; Best, R. B.; Mittal, J. Sequence Determinants of Protein Phase Behavior from a Coarse-Grained Model. PLOS Comput. Biol. 2018, 14 (1), e1005941. 10.1371/journal.pcbi.1005941.

(34) Tesei, G.; Schulze, T. K.; Crehuet, R.; Lindorff-Larsen, K. Accurate Model of Liquid-Liquid Phase Behavior of Intrinsically Disordered Proteins from Optimization of Single-Chain Properties. Proc. Natl. Acad. Sci. U. S. A. 2021, 118 (44), e2111696118. 10.1073/pnas.2111696118.

(35) Souza, P. C. T.; Alessandri, R.; Barnoud, J.; Thallmair, S.; Faustino, I.; Grünewald, F.; Patmanidis, I.; Abdizadeh, H.; Bruininks, B. M. H.; Wassenaar, T. A.; Kroon, P. C.; Melcr, J.; Nieto, V.; Corradi, V.; Khan, H. M.; Domański, J.; Javanainen, M.; Martinez-Seara, H.; Reuter, N.; Best, R. B.; Vattulainen, I.; Monticelli, L.; Periole, X.; Tieleman, D. P.; De Vries, A. H.; Marrink, S. J. Martini 3: A General Purpose Force Field for Coarse-Grained Molecular Dynamics. Nat. Methods 2021, 18 (4), 382–388. 10.1038/s41592-021-01098-3.

(36) Marrink, S. J.; Risselada, H. J.; Yefimov, S.; Tieleman, D. P.; De Vries, A. H. The MARTINI Force Field: Coarse Grained Model for Biomolecular Simulations. J. Phys. Chem. B 2007, 111 (27), 7812–7824. 10.1021/jp071097f.

(37) Marrink, S. J.; Monticelli, L.; Melo, M. N.; Alessandri, R.; Tieleman, D. P.; Souza, P. C. T. Two Decades of Martini: Better Beads, Broader Scope. WIREs Comput. Mol. Sci. 2023, 13 (1), e1620. 10.1002/wcms.1620.

(38) Benayad, Z.; Von Bülow, S.; Stelzl, L. S.; Hummer, G. Simulation of FUS Protein Condensates with an Adapted Coarse-Grained Model. J. Chem. Theory Comput. 2021, 17 (1), 525–537. 10.1021/acs.jctc.0c01064.

(39) Ingólfsson, H. I.; Rizuan, A.; Liu, X.; Mohanty, P.; Souza, P. C. T.; Marrink, S. J.; Bowers, M. T.; Mittal, J.; Berry, J. Multiscale Simulations Reveal TDP-43 Molecular-Level Interactions Driving Condensation. Biophys. J. 2023, 122 (22), 4370–4381. 10.1016/j.bpj.2023.10.016.

(40) Tsanai, M.; Frederix, P. W. J. M.; Schroer, C. F. E.; Souza, P. C. T.; Marrink, S. J. Coacervate Formation Studied by Explicit Solvent Coarse-Grain Molecular Dynamics with the Martini Model. Chem. Sci. 2021, 12 (24), 8521–8530. 10.1039/D1SC00374G.

(41) Zerze, G. H. Optimizing the Martini 3 Force Field Reveals the Effects of the Intricate Balance between Protein–Water Interaction Strength and Salt Concentration on Biomolecular Condensate Formation. J. Chem. Theory Comput. 2024, 20 (4), 1646–1655. 10.1021/acs.jctc.2c01273.

(42) Garg, A.; Brasnett, C.; Marrink, S. J.; Koren, K.; Kjaergaard, M. Oxygen Partitioning into Biomolecular Condensates Is Governed by Protein Density. May 5, 2024. 10.1101/2024.05.03.592328.

(43) Lescop, E.; Schanda, P.; Brutscher, B. A Set of BEST Triple-Resonance Experiments for Time-Optimized Protein Resonance Assignment. J. Magn. Reson. 2007, 187 (1), 163–169. 10.1016/j.jmr.2007.04.002.

(44) Wittekind, M.; Mueller, L. HNCACB, a High-Sensitivity 3D NMR Experiment to Correlate Amide-Proton and Nitrogen Resonances with the Alpha- and Beta-Carbon Resonances in Proteins. J. Magn. Reson. B 1993, 101 (2), 201–205. 10.1006/jmrb.1993.1033.

(45) Grzesiek, S.; Bax, A. Correlating Backbone Amide and Side Chain Resonances in Larger Proteins by Multiple Relayed Triple Resonance NMR. J. Am. Chem. Soc. 1992, 114 (16), 6291–6293. 10.1021/ja00042a003.

(46) Grzesiek, S.; Bax, A. Improved 3D Triple-Resonance NMR Techniques Applied to a 31 kDa Protein. J. Magn. Reson. 1969 1992, 96 (2), 432–440. 10.1016/0022-2364(92)90099-s.

(47) Lee, W.; Rahimi, M.; Lee, Y.; Chiu, A. POKY: A Software Suite for Multidimensional NMR and 3D Structure Calculation of Biomolecules. Bioinformatics 2021, 37 (18), 3041–3042. 10.1093/bioinformatics/btab180.

(48) Zhu, G.; Xia, Y.; Nicholson, L. K.; Sze, K. H. Protein Dynamics Measurements by TROSY-Based NMR Experiments. J. Magn. Reson. 2000, 143 (2), 423–426. 10.1006/jmre.2000.2022.

(49) Abraham, M. J.; Murtola, T.; Schulz, R.; Páll, S.; Smith, J. C.; Hess, B.; Lindahl, E. GROMACS: High Performance Molecular Simulations through Multi-Level Parallelism from Laptops to Supercomputers. SoftwareX 2015, 1–2, 19–25. 10.1016/j.softx.2015.06.001.

(50) Huang, J.; Rauscher, S.; Nawrocki, G.; Ran, T.; Feig, M.; De Groot, B. L.; Grubmüller, H.; MacKerell, A. D. CHARMM36m: An Improved Force Field for Folded and Intrinsically Disordered Proteins. Nat. Methods 2017, 14 (1), 71–73. 10.1038/nmeth.4067.

(51) Best, R. B.; Zhu, X.; Shim, J.; Lopes, P. E. M.; Mittal, J.; Feig, M.; MacKerell, A. D. Optimization of the Additive CHARMM All-Atom Protein Force Field Targeting Improved Sampling of the Backbone IZ, ψ and Side-Chain χ_1_ and χ_2_ Dihedral Angles. J. Chem. Theory Comput. 2012, 8(9), 3257–3273. 10.1021/ct300400x.

(52) Bernetti, M.; Bussi, G. Pressure Control Using Stochastic Cell Rescaling. J. Chem. Phys. 2020, 153 (11), 114107. 10.1063/5.0020514.

(53) Parrinello, M.; Rahman, A. Polymorphic Transitions in Single Crystals: A New Molecular Dynamics Method. J. Appl. Phys. 1981, 52 (12), 7182–7190. 10.1063/1.328693.

(54) Bussi, G.; Donadio, D.; Parrinello, M. Canonical Sampling through Velocity Rescaling. J. Chem. Phys. 2007, 126 (1), 014101. 10.1063/1.2408420.

(55) Jumper, J.; Evans, R.; Pritzel, A.; Green, T.; Figurnov, M.; Ronneberger, O.; Tunyasuvunakool, K.; Bates, R.; Žídek, A.; Potapenko, A.; Bridgland, A.; Meyer, C.; Kohl, S. A. A.; Ballard, A. J.; Cowie, A.; Romera-Paredes, B.; Nikolov, S.; Jain, R.; Adler, J.; Back, T.; Petersen, S.; Reiman, D.; Clancy, E.; Zielinski, M.; Steinegger, M.; Pacholska, M.; Berghammer, T.; Bodenstein, S.; Silver, D.; Vinyals, O.; Senior, A. W.; Kavukcuoglu, K.; Kohli, P.; Hassabis, D. Highly Accurate Protein Structure Prediction with AlphaFold. Nature 2021, 596 (7873), 583–589. 10.1038/s41586-021-03819-2.

(56) Souza, P. C. T.; Alessandri, R.; Barnoud, J.; Thallmair, S.; Faustino, I.; Grünewald, F.; Patmanidis, I.; Abdizadeh, H.; Bruininks, B. M. H.; Wassenaar, T. A.; Kroon, P. C.; Melcr, J.; Nieto, V.; Corradi, V.; Khan, H. M.; Domański, J.; Javanainen, M.; Martinez-Seara, H.; Reuter, N.; Best, R. B.; Vattulainen, I.; Monticelli, L.; Periole, X.; Tieleman, D. P.; De Vries, A. H.; Marrink, S. J. Martini 3: A General Purpose Force Field for Coarse-Grained Molecular Dynamics. Nat. Methods 2021, 18 (4), 382–388. 10.1038/s41592-021-01098-3.

(57) Kroon, P. C.; Grunewald, F.; Barnoud, J.; Van Tilburg, M.; Brasnett, C.; De Souza, P. C. T.; Wassenaar, T. A.; Marrink, S.-J. J. Martinize2 and Vermouth: Unified Framework for Topology Generation. June 23, 2025. 10.7554/eLife.90627.3.

(58) Souza, P. C. T.; Borges-Araújo, L.; Brasnett, C.; Moreira, R. A.; Grünewald, F.; Park, P.; Wang, L.; Razmazma, H.; Borges-Araújo, A. C.; Cofas-Vargas, L. F.; Monticelli, L.; Mera-Adasme, R.; Melo, M. N.; Wu, S.; Marrink, S. J.; Poma, A. B.; Thallmair, S. GōMartini 3: From Large Conformational Changes in Proteins to Environmental Bias Corrections. Nat. Commun. 2025, 16 (1), 4051. 10.1038/s41467-025-58719-0.

(59) Herzog, F. A.; Braun, L.; Schoen, I.; Vogel, V. Improved Side Chain Dynamics in MARTINI Simulations of Protein–Lipid Interfaces. J. Chem. Theory Comput. 2016, 12 (5), 2446–2458. 10.1021/acs.jctc.6b00122.

(60) De Jong, D. H.; Baoukina, S.; Ingólfsson, H. I.; Marrink, S. J. Martini Straight: Boosting Performance Using a Shorter Cutoff and GPUs. Comput. Phys. Commun. 2016, 199, 1–7. 10.1016/j.cpc.2015.09.014.

(61) Kim, H.; Fábián, B.; Hummer, G. Neighbor List Artifacts in Molecular Dynamics Simulations. J. Chem. Theory Comput. 2023, 19 (23), 8919–8929. 10.1021/acs.jctc.3c00777.

(62) Michaud-Agrawal, N.; Denning, E. J.; Woolf, T. B.; Beckstein, O. MDAnalysis: A Toolkit for the Analysis of Molecular Dynamics Simulations. J. Comput. Chem. 2011, 32 (10), 2319–2327. 10.1002/jcc.21787.

(63) Gowers, R.; Linke, M.; Barnoud, J.; Reddy, T.; Melo, M.; Seyler, S.; Domański, J.; Dotson, D.; Buchoux, S.; Kenney, I.; Beckstein, O. MDAnalysis: A Python Package for the Rapid Analysis of Molecular Dynamics Simulations; Austin, Texas, 2016; pp 98–105. 10.25080/Majora-629e541a-00e.

(64) Grünewald, F.; Marrink, S. J. Fgrunewald/Fast_forward: V0.0.1, 2022. 10.5281/ZENODO.6531921.

(65) Humphrey, W.; Dalke, A.; Schulten, K. VMD: Visual Molecular Dynamics. J. Mol. Graph. 1996, 14 (1), 33–38. 10.1016/0263-7855(96)00018-5.

(66) Brasnett, C.; Marrink, S. J. MartiniGlass: A Tool for Enabling Visualization of Coarse-Grained Martini Topologies. J. Chem. Inf. Model. 2025, 65 (7), 3137–3141. 10.1021/acs.jcim.4c02277.

(67) Banky, P.; Roy, M.; Newlon, M. G.; Morikis, D.; Haste, N. M.; Taylor, S. S.; Jennings, P. A. Related Protein– Protein Interaction Modules Present Drastically Different Surface Topographies Despite A Conserved Helical Platform. J. Mol. Biol. 2003, 330 (5), 1117–1129. 10.1016/S0022-2836(03)00552-7.

(68) Wang, J.; Choi, J.-M.; Holehouse, A. S.; Lee, H. O.; Zhang, X.; Jahnel, M.; Maharana, S.; Lemaitre, R.; Pozniakovsky, A.; Drechsel, D.; Poser, I.; Pappu, R. V.; Alberti, S.; Hyman, A. A. A Molecular Grammar Governing the Driving Forces for Phase Separation of Prion-like RNA Binding Proteins. Cell 2018, 174 (3), 688–699.e16. 10.1016/j.cell.2018.06.006.

(69) Ray, S.; Mason, T. O.; Boyens-Thiele, L.; Farzadfard, A.; Larsen, J. A.; Norrild, R. K.; Jahnke, N.; Buell, A. K. Mass Photometric Detection and Quantification of Nanoscale α-Synuclein Phase Separation. Nat. Chem. 2023, 15 (9), 1306–1316. 10.1038/s41557-023-01244-8.

(70) Burgers, P. P.; Bruystens, J.; Burnley, R. J.; Nikolaev, V. O.; Keshwani, M.; Wu, J.; Janssen, B. J. C.; Taylor, S. S.; Heck, A. J. R.; Scholten, A. Structure of smAKAP and Its Regulation by PKA-Mediated Phosphorylation. FEBS J. 2016, 283 (11), 2132–2148. 10.1111/febs.13726.

(71) Bugge, K.; Brakti, I.; Fernandes, C. B.; Dreier, J. E.; Lundsgaard, J. E.; Olsen, J. G.; Skriver, K.; Kragelund, B. B. Interactions by Disorder – A Matter of Context. Front. Mol. Biosci. 2020, 7, 110. 10.3389/fmolb.2020.00110.

(72) Sørensen, C. S.; Kjaergaard, M. Effective Concentrations Enforced by Intrinsically Disordered Linkers Are Governed by Polymer Physics. PNAS 2019, 116 (46), 23124–23131. 10.1101/577536.

(73) Prestel, A.; Wichmann, N.; Martins, J. M.; Marabini, R.; Kassem, N.; Broendum, S. S.; Otterlei, M.; Nielsen, O.; Willemoës, M.; Ploug, M.; Boomsma, W.; Kragelund, B. B. The PCNA Interaction Motifs Revisited: Thinking Outside the PIP-Box. Cell. Mol. Life Sci. 2019, 76 (24), 4923–4943. 10.1007/s00018-019-03150-0.

(74) Conicella, A. E.; Dignon, G. L.; Zerze, G. H.; Schmidt, H. B.; D’Ordine, A. M.; Kim, Y. C.; Rohatgi, R.; Ayala, Y. M.; Mittal, J.; Fawzi, N. L. TDP-43 α-Helical Structure Tunes Liquid–Liquid Phase Separation and Function. Proc. Natl. Acad. Sci. 2020, 117 (11), 5883–5894. 10.1073/pnas.1912055117.

(75) Brangwynne, C. P.; Tompa, P.; Pappu, R. V. Polymer Physics of Intracellular Phase Transitions. Nat. Phys. 2015, 11 (11), 899–904. 10.1038/nphys3532.

(76) Dass, R.; Mulder, F. A. A.; Nielsen, J. T. ODiNPred: Comprehensive Prediction of Protein Order and Disorder. Sci. Rep. 2020, 10 (1), 14780. 10.1038/s41598-020-71716-1.

