## Supplementary information for "Phase separation of protein kinase A regulatory subunits is driven by similar inter- and intra-molecular interactions involving the inhibitory segment"

**Contents:**

**1. Protein sequences**

**2. Supplementary figures:**

Fig. S1: Phase diagrams of RIα_1-63 and_ RIα_1-100_ as a function of crowder concentration and temperature.

Fig. S2: Backbone chemical shift assignment of RIα_1-100_ as a function of temperature and pH.

Fig. S3: Backbone chemical shift assignment of RIα_1-63_.

Fig. S4: Phase diagram of RIα_1-93,_ RIα_1-100_(6R-A) and RIα_1-100_(6K-A)

Fig. S5: Slab setup for simulation of condensate

**1. Protein sequences:**

All proteins were expressed in pet28b using NdeI and XhoI cloning sites and have the following N-terminal sequence from the plasmid, which is removed by TEV cleavage:

MGSSHHHHHHSSGLVPRGSHM

TEV cleavage sites are highlighted in yellow, where | indicates the scissile peptide bond.

Point mutations are highlighted in red.

**RIα_1-63_ (WT):**

ENLYFQ|SMESGSTAASEEARSLRECELYVQKHNIQALLKDSIVQLCTARPERPMAFLREYFERLEKEEAK

**RIα_1-93_ (WT):**

ENLYFQ|SMESGSTAASEEARSLRECELYVQKHNIQALLKDSIVQLCTARPERPMAFLREYFERLEKEEAKQIQNLQKAGTRTDSREDEISPPPPNPVVKG

**RIα_1-100_ (WT):**

ENLYFQ|SMESGSTAASEEARSLRECELYVQKHNIQALLKDSIVQLCTARPERPMAFLREYFERLEKEEAKQIQNLQKAGTRTDSREDEISPPPPNPVVKGRRRRGAI

**RIα_1-100_ (6RtoK):**

ENLYFQ|SMESGSTAASEEARSLRECELYVQKHNIQALLKDSIVQLCTARPERPMAFLREYFERLEKEEAKQIQNLQKAGTKTDSKEDEISPPPPNPVVKGKKKKGAI

**RIα_1-100_ (6RtoA):**

ENLYFQ|SMESGSTAASEEARSLRECELYVQKHNIQALLKDSIVQLCTARPERPMAFLREYFERLEKEEAKQIQNLQKAGTATDSAEDEISPPPPNPVVKGAAAAGAI

**RIα_64-105_ (WT):**

MESGSTAASEEARSLRECELYVQKHNIQALLKDSIVQLCTARPERPMAFLREYFERLEKEEAKENLYFQ|SQIQNLQKAGTRTDSREDEISPPPPNPVVKGRRRRGAISAEVY

**2. Supplementary figures**

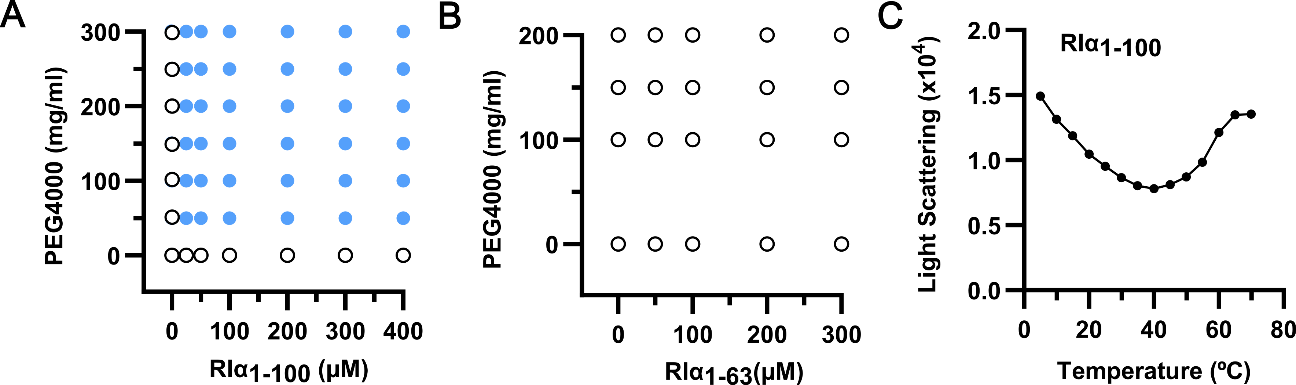

**Figure S1**: **Phase diagrams of** **RIα_1-100_ and RIα_1-63_ as a function of crowder concentration and temperature.** Representative *in vitro* phase diagram of **A)** RIα_1-100_ wild type and **B)** RIα_1-63_ mutant at varying concentration of protein and crowder generated using light scattering data and/or imaged DIC microscopy. **C)** Temperature dependent phase separation of RIα_1-100_ observed by static light scattering at 100 µM protein concentration in the presence of 100 mg/ml PEG4000 and 50 mM KCl.

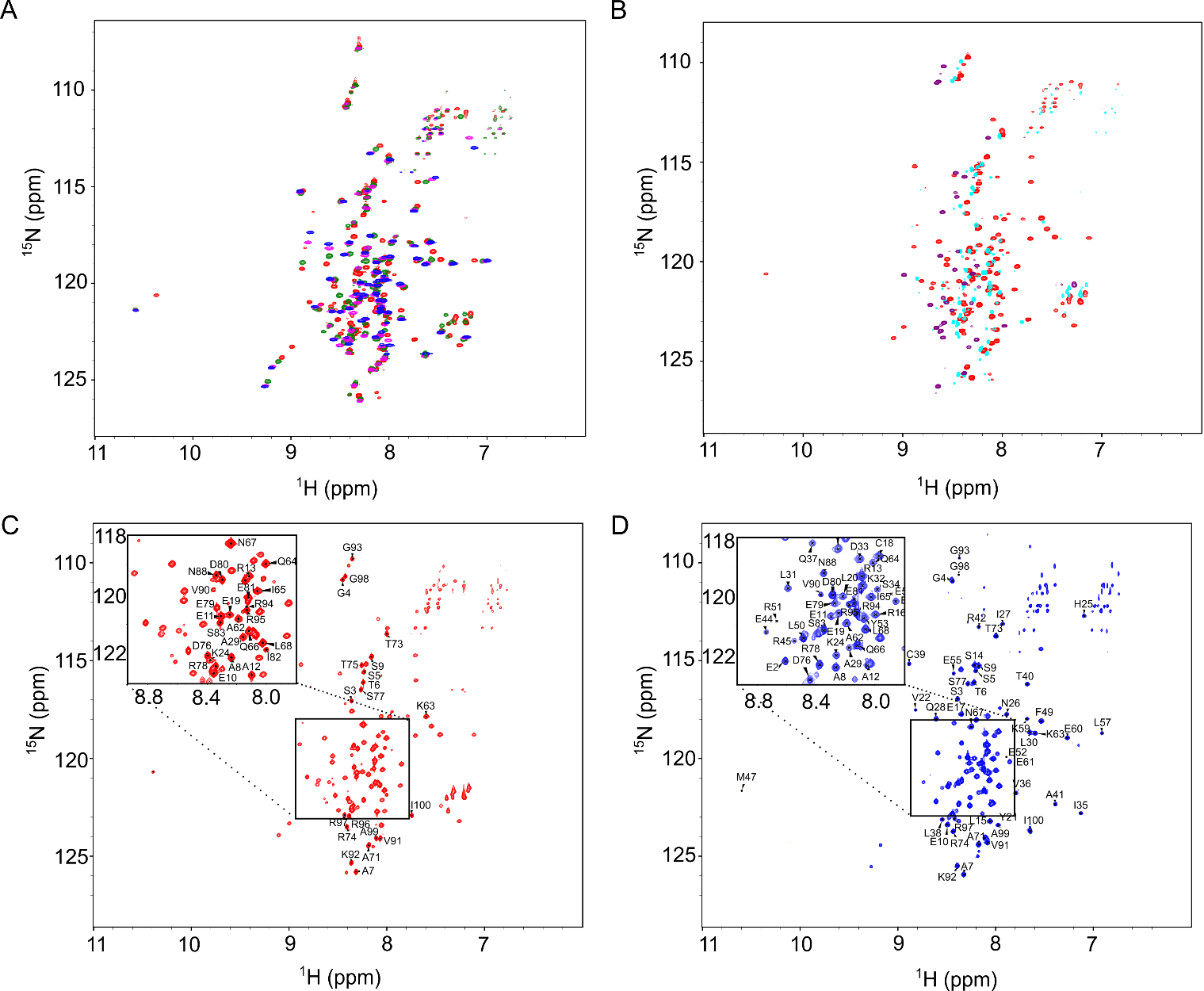

**Figure S2: Backbone chemical shift assignment of RIα_1-100_ as a function of temperature and pH.** A) Overlaid TROSY-based ^15^N-^1^H HSQC spectra of RIα_1-100_, recorded at 310 K and pH 4 (red), pH 5 (green), pH 6 (magenta) and pH 7 (blue). B) Overlaid TROSY-based ^15^N-^1^H HSQC spectra of RIα_1-100_  , recorded at pH 4 at 278 K (purple), 298 K (turquoise) and 310 K (Red). C) TROSY-based ^15^N-^1^H HSQC spectrum of RIα_1-100_ at pH 4 and 310 K, with assignments for residues from the N-terminal and C-terminal IDR tail. D) TROSY-based ^15^N-^1^H HSQC spectrum of RIα_1-100_ at pH 7 and 298 K, showing assignments for residues from both the folded D/D domain and the IDR.

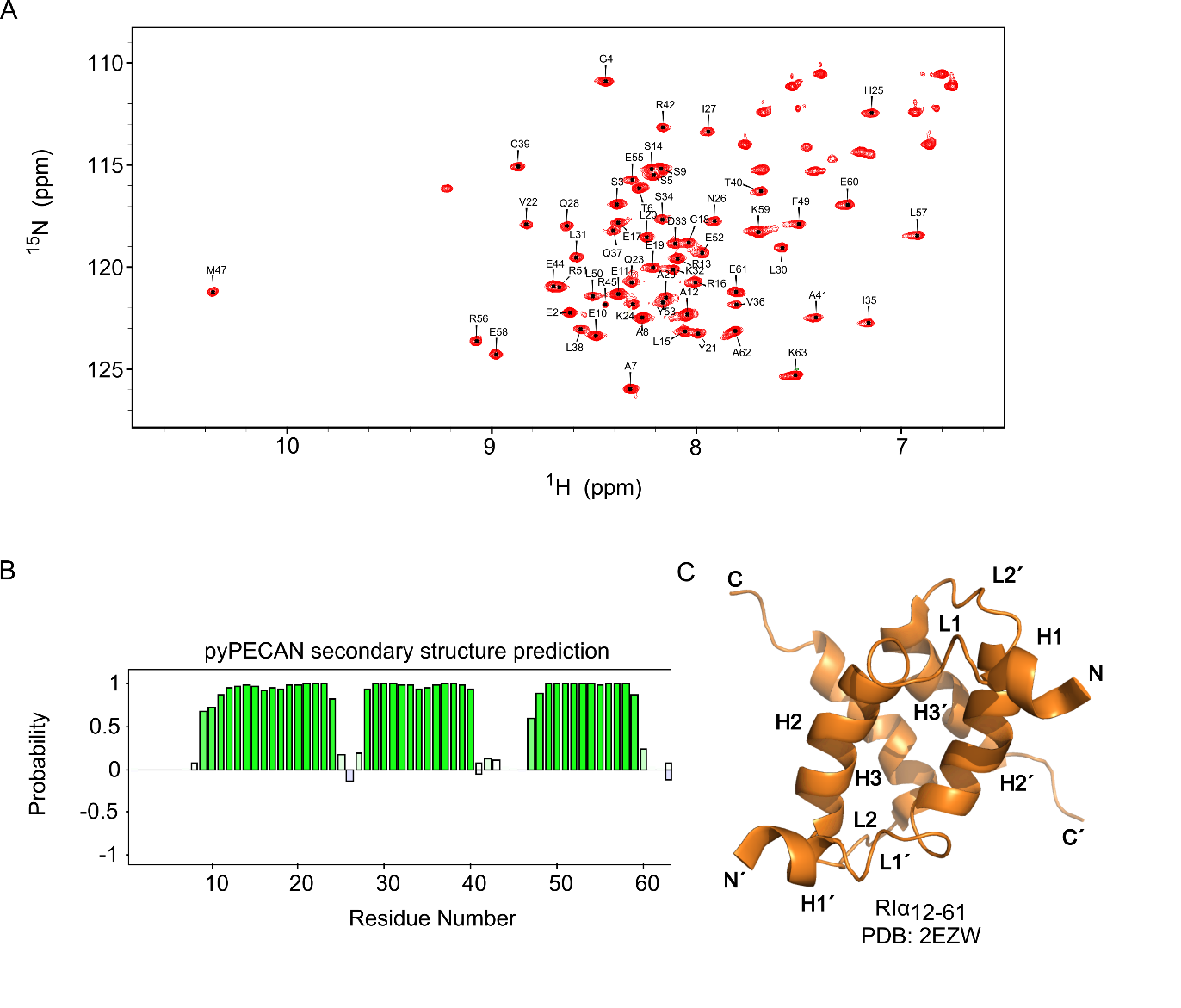

**Figure S3: Backbone chemical shift assignment of** **RIα_1-63_.** A) Assigned 2D ^15^N-^1^H HSQC spectrum of RIα_1-63_ at pH 7 and 298 K. B) Secondary structure prediction of RIα_1-63_ based on NMR secondary chemical shifts. C) Published NMR structure of the folded D/D domain (RIα_12-61_, PDB: 2EZW).

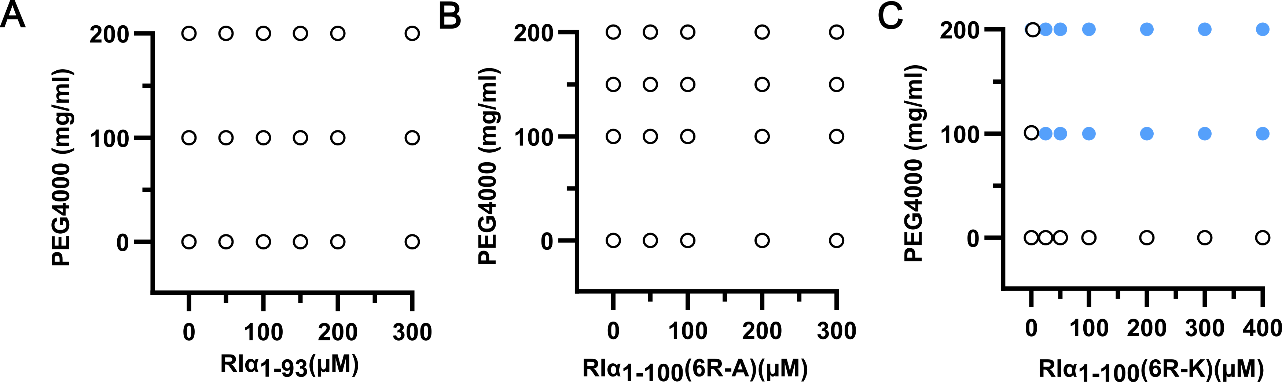

**Figure S4: Phase diagram of RIα_1-93,_ RIα_1-100_(6R-A) and RIα_1-100_(6R-K)**. Representative *in vitro* phase diagram of A) RIα_1-93_, B) RIα_1-100_(6R-A) and C) RIα_1-100_(6R-K) mutants with increasing concentration of protein and crowder. Data recorded as in Fig. S1.

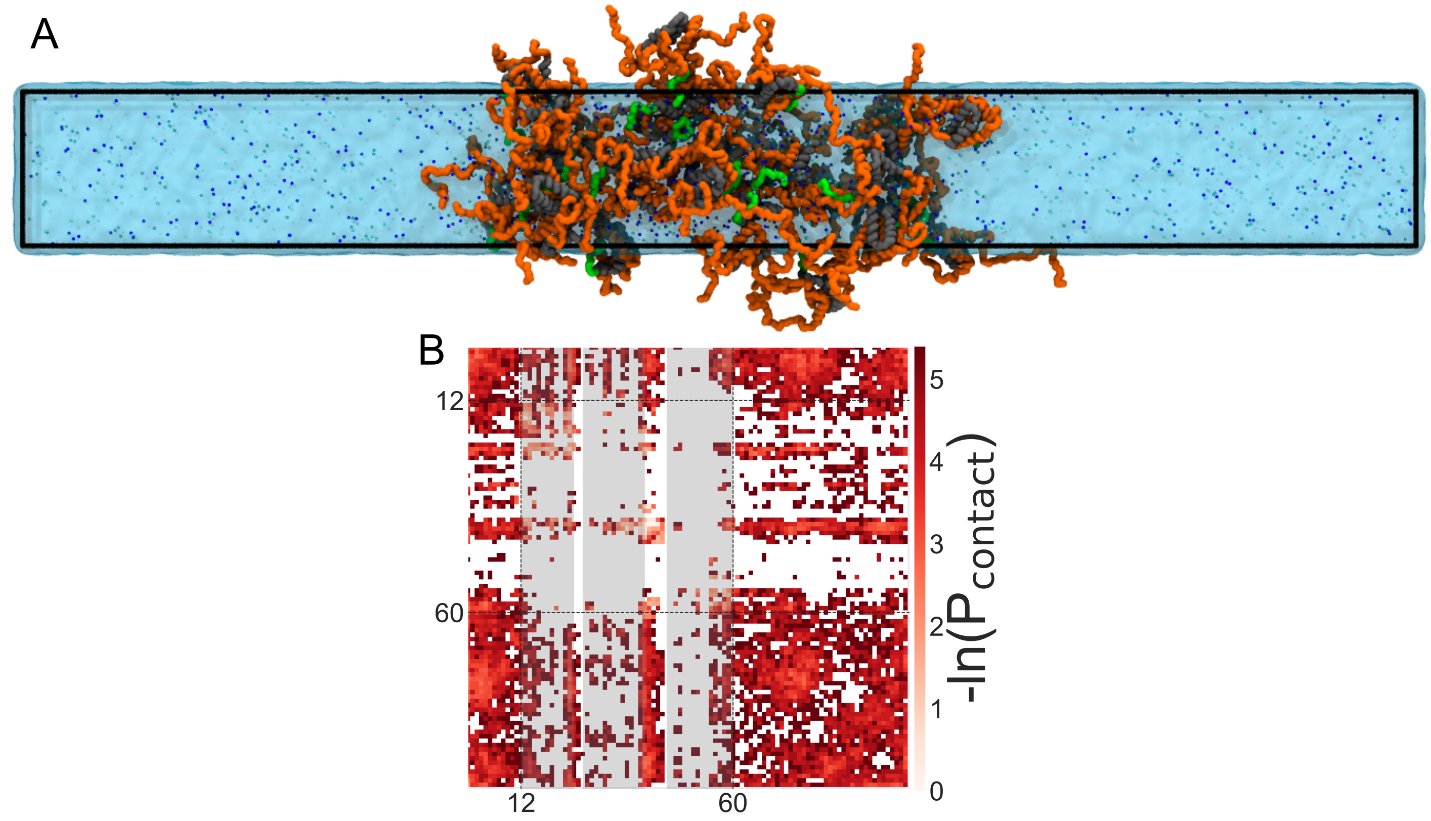

**Figure S5: Slab setup for simulation of condensate.** A) Beginning from a slab condensate configuration, the PKA RIα_1-100_ is simulated for 10 μs, and the intermolecular contacts are calculated. B) The contacts between molecules show identical pattern prevalences to simulations of self-associating condensates.
